# The *Chlamydia* protein CpoS modulates the inclusion microenvironment and restricts the interferon response by acting on Rab35

**DOI:** 10.1101/2022.02.18.481055

**Authors:** Karsten Meier, Lana H. Jachmann, Lucía Pérez, Oliver Kepp, Raphael H. Valdivia, Guido Kroemer, Barbara S. Sixt

**Author notes:** Senior authors (Raphael H. Valdivia, Guido Kroemer, Barbara S. Sixt).

## Abstract

The obligate intracellular bacterium *Chlamydia trachomatis* inserts into the membrane of its vacuole (the inclusion) a family of poorly characterized Inc proteins. While the Inc CpoS was recently revealed as a critical suppressor of host cellular immune surveillance, the underlying mechanism remained unknown. By complementing a *cpoS* mutant with modified variants of CpoS, we found that CpoS blocks distinct cellular defense responses through distinct mechanisms. Specifically, we show that the ability of CpoS to interact with Rab GTPases is not only instrumental to its ability to mediate lipid transport to the inclusion, but also key to CpoS-mediated inhibition of type I interferon responses. Indeed, depletion of Rab35 can phenocopy the respective defect of the *cpoS* mutant. Unexpectedly, we found that CpoS is also essential for the formation of inclusion microdomains that control the spatial organization of multiple Incs involved in signaling and modulation of the host cellular cytoskeleton. Overall, our findings highlight the modulation of membrane trafficking as a pathogenic immune evasion strategy and the role of Inc-Inc interactions in shaping the inclusion microenvironment.

## INTRODUCTION

Intracellular bacteria often thrive within membrane-bound bacteria-containing vacuoles that provide an adequate growth niche and protection from host cell-autonomous immune surveillance. The vacuole membrane, which serves as a hub for interactions with host proteins, is heavily modified by bacterial factors. A mechanistic understanding of how these virulence factors maintain the function of the pathogenic compartment could help identify novel targets for therapeutic interventions.

A prominent example of bacteria that excel as manipulators of mammalian cell biology are the obligate intracellular pathogens of the genus *Chlamydia*, which are responsible for a range of diseases in humans and animals (Kuo et al., 2011, Longbottom et al., 2003). For instance, *Chlamydia trachomatis*, a major human pathogen, is the causative agent of blinding trachoma (serovars A-C) (Taylor et al., 2014), and a prevalent agent of urogenital infections that can cause infertility and pregnancy complications (serovars D-K) (Mylonas, 2012). Moreover, *C. trachomatis* can cause invasive diseases like lymphogranuloma venereum (LGV, serovars L1-L3) (Ceovic et al., 2015).

*Chlamydia* spp. are characterized by a biphasic development (Ward, 1988). The *Chlamydia* vacuole, termed inclusion, is established when the elementary body (EB), the infectious non-replicative form of the pathogen, invades a host cell. The EB then differentiates into the reticulate body (RB), the non-infectious replicative form, which multiplies inside the inclusion. Eventually, RBs differentiate back into EBs, which are released by host cell lysis or extrusion, to infect other cells. In cultured cells infected with *C. trachomatis*, this cycle is typically completed within 48 to 72 hours (Hybiske et al., 2007).

*Chlamydia* spp. modify the membrane of their vacuole by secreting a class of effector proteins (Incs) that are inserted into the inclusion membrane (Bannantine et al., 2000). Incs interact extensively with host proteins and with each other (Gauliard et al., 2015, Mirrashidi et al., 2015), thus mediating a range of processes, such as homotypic fusion of inclusions (Hackstadt et al., 1999), formation of ER-inclusion contact sites (Derre et al., 2011), cytoskeletal rearrangements (Dumoux et al., 2015, Haines et al., 2021, Kokes et al., 2015), and modulation of host vesicular transport (Delevoye et al., 2008, Rzomp et al., 2006). The high number of Incs encoded by *Chlamydia* genomes, over 50 in *C. trachomatis* (> 5% of its protein-coding genes) (Weber et al., 2015), highlights their biological importance. Moreover, the repertoire and sequence composition of Incs varies between *Chlamydia* species and among biovars of *C. trachomatis* (Almeida et al., 2012, Lutter et al., 2012), suggesting Incs could be determinants of host and tissue tropism.

One of these polymorphic Incs is CpoS (also known as CTL0481 in *C. trachomatis* serovar L2 and CT229 in serovar D, henceforth referred to as CpoS(L2) and CpoS(D)). Exploiting the novel tools for genetic manipulation of *Chlamydia* spp. (Sixt et al., 2016), we and others recently discovered that CpoS-deficient *C. trachomatis* L2 strains induce a robust STING-dependent type I interferon (IFN) response and rapid necrotic and apoptotic death in infected cells (Sixt et al., 2017, Weber et al., 2017). CpoS-deficient bacteria are also impaired in their ability to produce infectious EBs and to survive in the murine genital tract (Sixt et al., 2017, Weber et al., 2017). Together, these findings revealed CpoS as a key virulence factor that suppresses cell-autonomous immunity. While CpoS(L2) and CpoS(D) can interact with host Rab GTPases (Faris et al., 2019, Mirrashidi et al., 2015, Rzomp et al., 2006, Sixt et al., 2017) and with other *Chlamydia* Incs (Gauliard et al., 2015, Sixt et al., 2017, Spaeth et al., 2009), the significance of these interactions in suppressing cellular defense pathways remained elusive.

To determine the mechanism by which CpoS subverts the host cellular defense, we analyzed the capacity of distinct variants of CpoS, including orthologous proteins and modified variants, to complement the phenotypes observed in a *cpoS* null mutant and to restore interactions with host and *Chlamydia* proteins. We found that all tested CpoS orthologs restored the full repertoire of tested phenotypes and interactions. We further provide evidence that CpoS dampens cell death and IFN responses through distinct molecular interactions and that the inhibition of IFN responses is linked to the CpoS-mediated recruitment of Rab GTPases to the inclusion. Unexpectedly, we also uncovered a role of CpoS in the establishment and/or maintenance of inclusion membrane microdomains, highlighting the significance of Inc-Inc associations in shaping *Chlamydia*-host interactions.

## RESULTS

### CpoS orthologs broadly complement a *cpoS* null mutant in *C. trachomatis* L2

Proteins orthologous to CpoS can be found in all serovars of *C. trachomatis*, yet their sequences vary (Fig 1A) and cluster according to disease tropism (i.e. ocular, urogenital, or LGV) (Almeida et al., 2012, Lutter et al., 2012). CpoS orthologs are also found in the mouse pathogen *C. muridarum* and in the swine pathogen *C. suis*, but their amino acid sequence similarities to CpoS in *C. trachomatis* are relatively low (Fig 1A). We thus tested if the function of CpoS in blocking cell-autonomous defenses is conserved across serovars and species.

**Figure 1.**
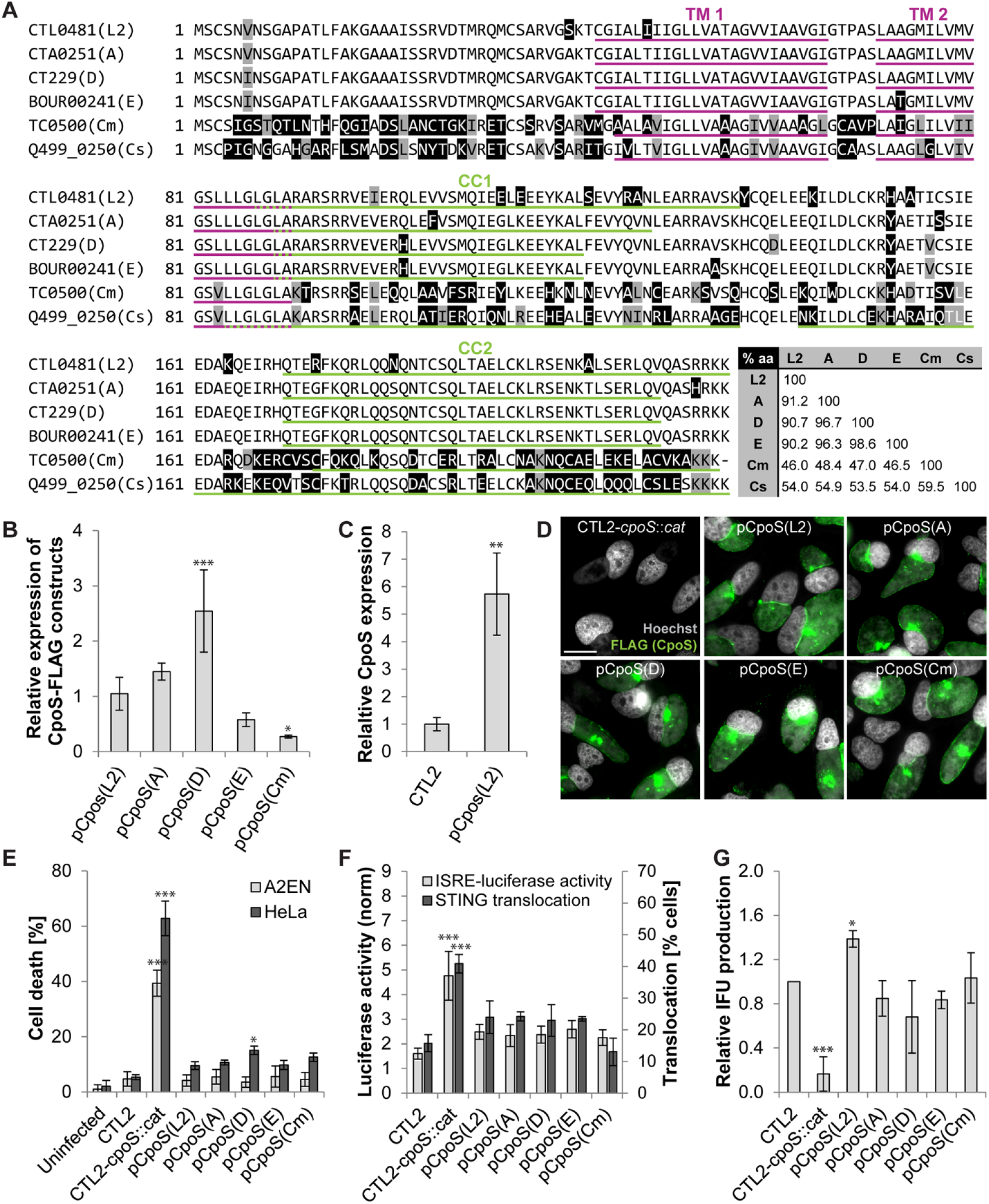
CpoS orthologs broadly complement a *cpoS* null mutant in *C. trachomatis* L2. **(A)** Protein sequence alignment and matrix of pairwise identities for selected orthologs (CTL0481, *C. trachomatis* L2/434/Bu; CTA0251, *C. trachomatis* A/HAR-13; CT229, *C. trachomatis* D/UW- 3/CX; BOUR00241, *C. trachomatis* E/Bour; TC0500, *C. muridarum* Nigg; Q499_0250, *C. suis* MD56; magenta, predicted transmembrane (TM) domains; green, predicted coiled-coil (CC) motifs). **(B-C)** Relative expression levels of CpoS orthologs in infected HeLa cells (MOI20, 32 hpi) determined by western blot analysis using anti-FLAG (B) or anti-CpoS (C) antibodies. **(D)** Localization of CpoS orthologs in infected HeLa cells (MOI5, 22 hpi, scale=20 µm). **(E)** Ability of CpoS orthologs to block cell death in infected cells (MOI10, 24 hpi). **(F)** Ability of CpoS orthologs to dampen the IFN response. Left y-axis: Luciferase activity in infected A2EN-ISRE reporter cells (MOI10, 14 hpi). Right y-axis: Frequency of STING translocation in MLFs (MOI5, 18 hpi, at least 270 cells per condition analyzed). **(G)** Ability of CpoS orthologs to restore IFU production in HeLa cells (measured at 36 hpi). All quantitative data in Fig 1 represent mean±SD [n=6 (A2EN in (E)), n=3 (else); student t test (C) or one-way ANOVA (else): indicated are significant differences compared to pCpoS(L2) (B) or CTL2 (else)].

We first disrupted *cpoS* in *C. trachomatis* L2 (CTL2) with an insertion carrying a *cat* resistance gene (Fig S1A). This strain, CTL2-*cpoS*::*cat*, like the previously described strain CTL2-*cpoS*::*bla* (Sixt et al., 2017), does not express any functional full-length CpoS protein. We then complemented the CTL2-*cpoS*::*cat* mutant with plasmids driving the expression of CpoS(L2) or CpoS orthologs derived from other *C. trachomatis* serovars [CpoS(A), CpoS(D), CpoS(E)] or *C. muridarum* [CpoS(Cm)], placed under the control of their native promoters (Fig 1A and Table S1). All orthologs were tagged with a C-terminal FLAG epitope, a modification that does not compromise the function of CpoS(L2) (Sixt et al., 2017). Immunoblots of infected human cervical epithelial (HeLa) cells indicated that the CpoS orthologs were expressed at variable amounts (Fig 1B and S1B), though always above the endogenous level of CpoS in CTL2 (compare Fig 1B and 1C). Moreover, immunofluorescence microscopy confirmed that all CpoS orthologs correctly localized to the inclusion membrane in infected cells (Fig 1D).

Consistent with former observations for CpoS-deficient strains (Sixt et al., 2017), infection with CTL2-*cpoS*::*cat,* but not wild-type CTL2, induced the premature death of HeLa and human endocervical epithelial (A2EN) cells, as indicated by the release of lactate dehydrogenase (LDH) detectable at 24 hours post infection (hpi) (Fig 1E). Prior to the induction of cell death, at 14 hpi with CTL2-*cpoS*::*cat*, we detected an enhanced expression of luciferase in an A2EN reporter cell line, which expresses luciferase under the control of the IFN-stimulated response element (ISRE) (Fig 1F). This finding indicates enhanced secretion of type I IFNs in cultures infected with the CpoS-deficient mutant. Moreover, when we infected *Goldenticket* mouse lung fibroblasts (MLFs) that express HA-tagged STING, we observed that at 18 hpi the frequency of cells displaying STING activation (defined as microscopically detectable STING translocation from the ER to perinuclear vesicles) was increased in cultures infected with CTL2-*cpoS*::*cat* (Fig 1F). Plasmid-driven reconstitution of CpoS(L2) into CTL2-*cpoS*::*cat* abolished host cell death induction (Fig 1E) and reduced STING translocation and induction of IFN-driven luciferase production (Fig 1F). Of note, expression of each of the tested CpoS orthologs, including the more distantly related ortholog CpoS(Cm), in CTL2-*cpoS*::*cat*, protected infected cells from premature death and blunted the activation of the IFN response (Fig 1E-F).

We next tested if the different CpoS orthologs restored the loss of replication potential observed in *cpoS* null mutants (Sixt et al., 2017, Weber et al., 2017). HeLa cultures were infected and the presence of infectious progeny was assessed at 36 hpi by quantifying the number of inclusion forming units (IFUs, i.e. EBs) in cell lysates. CTL2-*cpoS*::*cat* had a strongly reduced ability to form infectious progeny (Fig 1G). Complementation with CpoS(L2), or any of the tested CpoS orthologs, restored IFU production to normal levels (Fig 1G).

Taken together, these findings indicate that CpoS orthologs, despite their variance in amino acid homology, had conserved activities in protecting cells from death, restricting IFN responses, and restoring the full replication potential of *Chlamydia*.

### CpoS variants with altered coiled-coil motifs display differential abilities to counteract cell-autonomous defenses

CpoS(L2) possesses a short N-terminal domain (amino acids (aa) 1-41) that contains the signal sequence for type III secretion (Subtil et al., 2005), a hydrophobic domain (aa 42-90) with two transmembrane regions that mediate the insertion into the inclusion membrane, as well as a longer C-terminal domain (aa 91-215) of unknown function (Fig 2A). The C-terminal domain is predicted to be exposed to the host cell cytosol during infection and to contain two coiled-coil (CC) motifs, which are located approximately at aa 87-136 (CC1) and aa 170-215 (CC2) (Fig 2A). CpoS orthologs display a similar topology and domain structure (Fig 1A).

**Figure 2.**
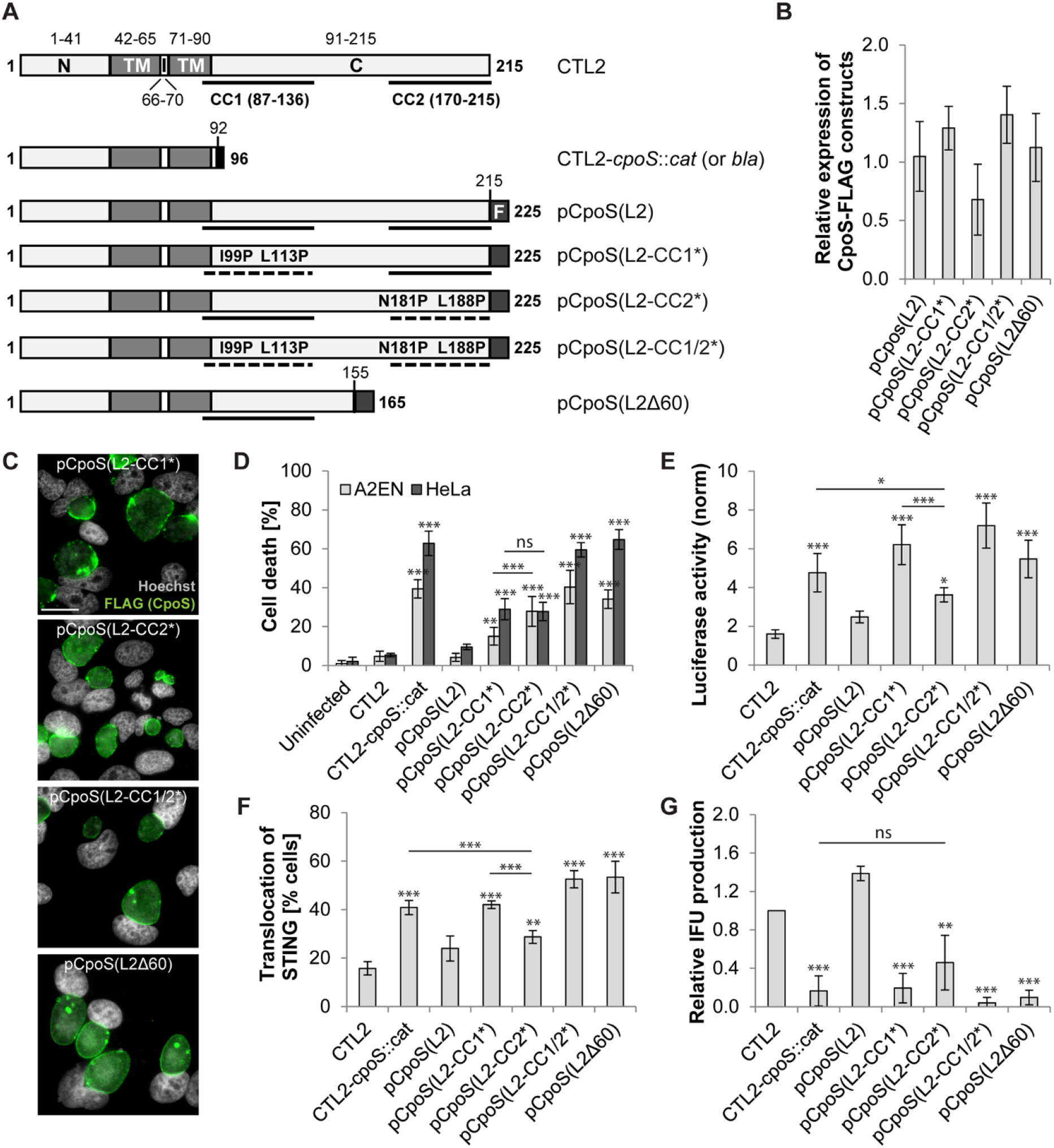
CpoS variants with altered coiled-coil motifs display differential abilities to counteract cell-autonomous defenses. **(A)** Predicted domain structure of CpoS(L2) expressed by wild-type, mutant, and complemented strains [N-terminal (N), transmembrane (TM), intra-inclusion (I), and C-terminal (C) domains; coiled-coil (CC) motifs; FLAG (F) tag]. **(B)** Relative expression levels of CpoS variants in infected HeLa cells (MOI20, 32 hpi) determined by western blot analysis using anti-FLAG antibodies. **(C)** Localization of CpoS variants in infected HeLa cells (MOI5, 22 hpi, scale=20 µm). **(D-G)** Ability of CpoS variants to prevent cell death (D), block ISRE-driven luciferase expression (E), block STING translocation (F), and restore IFU production (G). Details as described in Fig 1E-G. Note, the experiments were conducted together with those shown in Fig 1E-G and share the same controls. All quantitative results in Fig 2 represent mean±SD [n=6 (A2EN in (D)), n=3 (else); one-way ANOVA: indicated are significant differences compared to pCpoS(L2) (B) or CTL2 (else, unless indicated otherwise)].

To determine the contribution of the CC motifs to CpoS functions, we transformed CTL2-*cpoS*::*cat* with plasmids driving expression of various FLAG-tagged variants of CpoS(L2) under the control of the CpoS(L2) promoter. These variants included unmodified CpoS(L2), a truncated CpoS(L2) lacking 60 aa (including CC2) at the C-terminus [CpoS(L2Δ60)], and CpoS(L2) variants in which the α-helices constituting either CC1 or CC2 or both were disrupted by aa substitutions introducing proline residues [CpoS(L2-CC1*), CpoS(L2-CC2*), CpoS(L2-CC1/2*)] (Fig 2A and Table S1). Western blot and immunofluorescence experiments confirmed that all variants were expressed at comparable levels and localized to the inclusion membrane in infected HeLa cells (Fig 2B-C and S1B).

Expression of CpoS(L2Δ60) or CpoS(L2-CC1/2*) in CTL2-*cpoS*::*cat* failed to protect infected cells from cell death (Fig 2D) and resulted in an exaggerated IFN response (Fig 2E-F). Interestingly, while expression of CpoS(L2-CC1*) or CpoS(L2-CC2*) partially protected cells from death (Fig 2D), only CpoS(L2-CC2*) reduced STING activation and IFN-driven luciferase production in the reporter cells (Fig 2E-F). CpoS(L2-CC2*) was also the only variant tested that seemed to partially restore IFU production (Fig 2G).

Taken together, our findings demonstrate that both suppression of cell death and dampening of IFN production critically rely on the C-terminal domain of CpoS, but through distinct mechanisms. We speculated that specific mutants in this C-terminal domain offer the opportunity to identify the molecular basis for CpoS-mediated suppression of the IFN response.

### CpoS deficiency disrupts formation of inclusion membrane microdomains

*Chlamydia* Incs interact with each other, presumably to form higher order interaction networks. CpoS(D) was shown to interact with the Inc CT223 (IPAM, inclusion protein acting on microtubules) in a yeast two-hybrid system (Spaeth et al., 2009) and with the Incs IPAM, CT115 (IncD), and CT222 in a bacterial two-hybrid system (Gauliard et al., 2015). Among these possible interactors, only IPAM (also known as CTL0476 in *C. trachomatis* L2) was found to co-immunoprecipitate with CpoS(L2)-FLAG (Sixt et al., 2017).

We confirmed the interaction between CpoS(L2) and IPAM(L2) by co-immunoprecipitation (Co-IP) using lysates of HeLa cells that had been co-infected with two strains, one expressing CpoS(L2)-FLAG and the other IPAM(L2)-MYC. Because the inclusions inhabited by the two strains fuse, due to the action of IncA (Hackstadt et al., 1999), this system brought together CpoS(L2)-FLAG and IPAM(L2)-MYC in the same inclusion membrane. Thus, when CpoS(L2)- FLAG was immunoprecipitated using anti-FLAG beads, immunoblots confirmed Co-IP of IPAM(L2)-MYC (Fig 3A).

**Figure 3.**
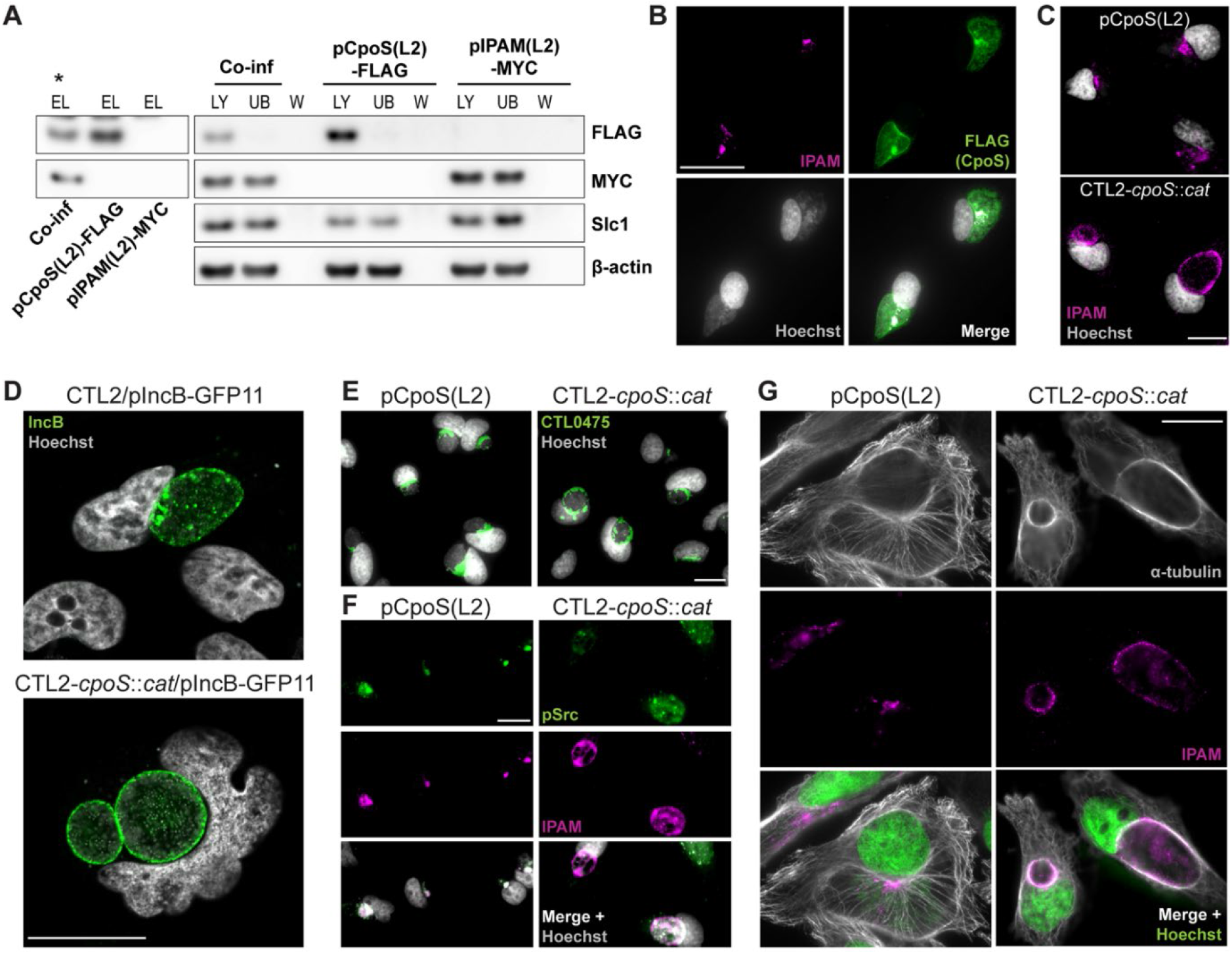
CpoS deficiency disrupts formation of inclusion membrane microdomains. **(A)** Co-IP confirms interaction between CpoS and IPAM. HeLa cells were co-infected with CTL2/pIPAM(L2)-MYC and CTL2-*cpoS*::*cat*/pCpoS(L2)-FLAG or were infected with each strain individually (MOI10). CpoS-FLAG was precipitated from lysates at 26 hpi. Samples were analyzed by western blot analysis (LY, lysate; UB, unbound; W, wash; EL, eluate; * indicates visible Co-IP). **(B)** Enrichment of CpoS-FLAG in IPAM-containing microdomains in HeLa cells infected with CTL2-*cpoS*::*cat*/pCpoS(L2)-FLAG (MOI5, 24 hpi, scale=40 µm).**(C-F)** Localization of IPAM (C), IncB (D), CTL0475 (E) and active Src kinases (F) in infected HeLa cells (MOI5-10, 22-24 hpi, scale=20 µm). The proteins were detected by immunostaining of endogenous proteins (C, E-F) or by using the split-GFP approach (D). **(G)** Architecture of the MT cytoskeleton in infected HeLa cells (MOI5, 22 hpi, scale=20 µm).

IPAM expressed endogenously by CTL2 is known to display a non-uniform distribution at the inclusion membrane with a pronounced accumulation at discrete regions termed inclusion membrane microdomains (Alzhanov et al., 2009, Bannantine et al., 2000, Dumoux et al., 2015, Weber et al., 2015). By immunofluorescence microscopy, we observed that CpoS was particularly abundant at these microdomains (Fig 3B). Strikingly, in cells infected with CTL2-*cpoS*::*cat*, IPAM localization was no longer restricted to inclusion microdomains (Fig 3C).

Several other Incs have been reported to accumulate at microdomains (Mital et al., 2010, Weber et al., 2015), including CTL0484 (IncB) and CTL0475, the *C. trachomatis* L2 ortholog of CT222, which was shown to interact with CpoS(D) and IPAM(D) in a bacterial two-hybrid assay (Gauliard et al., 2015). Like IPAM, IncB and CTL0475 displayed a redistribution away from microdomains during infection with the *cpoS* mutant (Fig 3D-E). The same was observed for activated (phosphorylated) host Src kinases (Fig 3F), which are typically also recruited to the inclusion microdomains (Mital et al., 2010, Weber et al., 2015).

Although the function of inclusion microdomains is not well understood, IPAM has been proposed to modulate the host microtubule (MT) network and cytokinesis by recruiting centrosomes to the proximity of the microdomains (Dumoux et al., 2015). Accordingly, we observed that the loss of CpoS affected MT architecture in infected cells. During infection with CTL2, MT-organizing centers were typically found in close apposition to IPAM-positive inclusion membrane patches, and the inclusions were surrounded by a “cage” of MTs, as previously described (Wesolowski et al., 2017). Of note, in the absence of CpoS, this accumulation of MTs at the periphery of inclusions was more pronounced (Fig 3G).

Altogether, these data demonstrate that CpoS deficiency leads to a general dispersal of inclusion microdomains or even prevents their formation, suggesting that the loss of a single Inc can have significant effects on the entire Inc interaction network.

### The disruption of inclusion microdomains is not linked to an augmented cell-autonomous defense

To explore a potential link between inclusion microdomains and the subversion of cell-autonomous defenses by CpoS, we evaluated the capacity of CpoS orthologs and variants to mediate interactions with IPAM and to restore microdomain formation.

Each of the tested FLAG-tagged CpoS orthologs co-precipitated with IPAM(L2)-MYC during immunoprecipitation using anti-FLAG beads (Fig 4A). Interestingly, CpoS(L2-CC1*) was the only modified CpoS(L2) variant that still interacted with IPAM (Fig 4A). Moreover, immunofluorescence microscopy showed that the formation of IPAM microdomains was restored by expression of each of the CpoS orthologs and by CpoS(L2-CC1*), but not by the variants that failed to interact with IPAM (Fig 4B).

**Figure 4.**
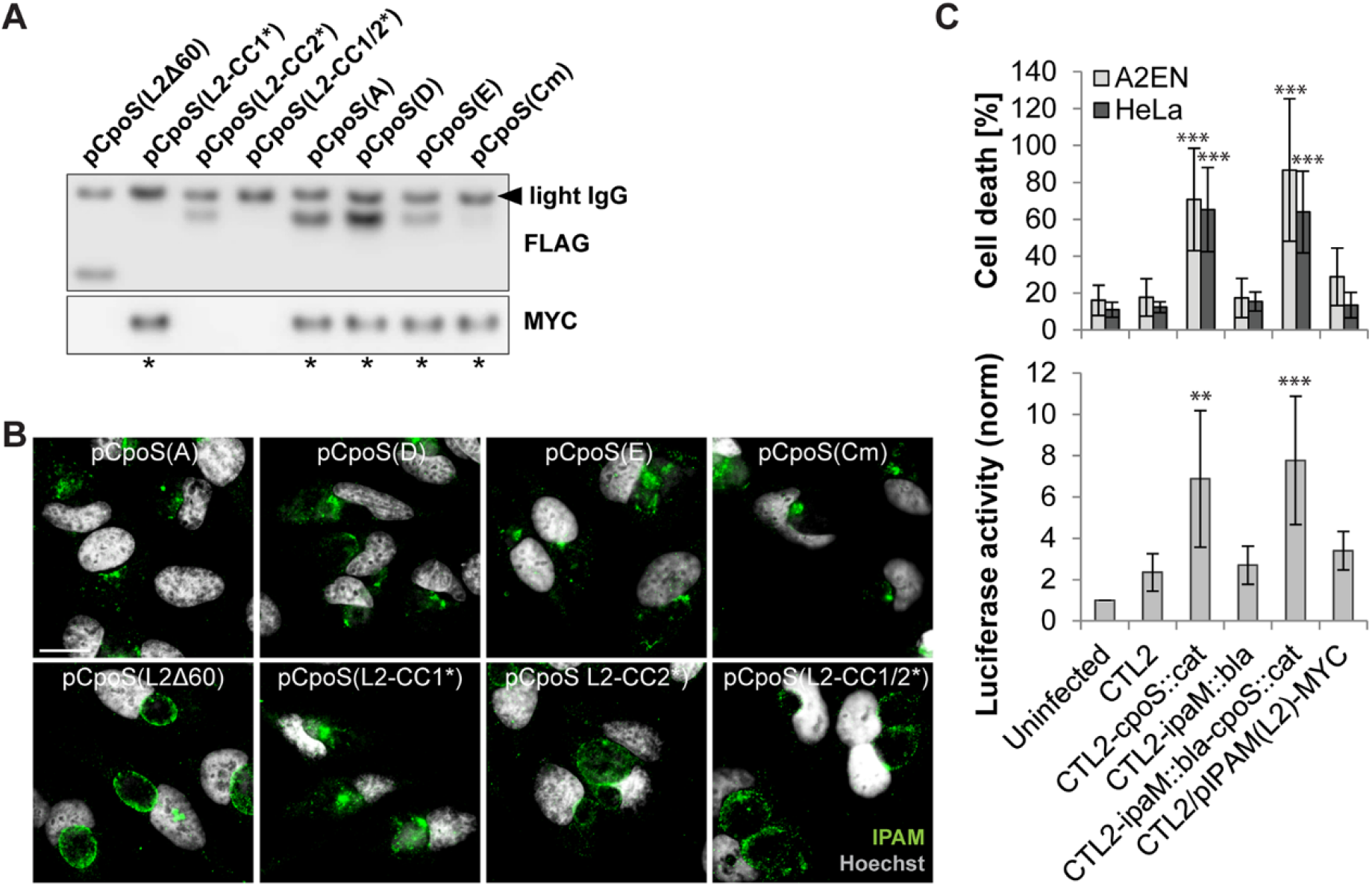
The disruption of inclusion microdomains is not linked to an augmented cell-autonomous defense. **(A)** Co-IP reveals a differential ability of CpoS variants to interact with IPAM. HeLa cells were co-infected with CTL2/pIPAM(L2)-MYC and derivatives of CTL2-*cpoS*::*cat* expressing variants of CpoS-FLAG (MOI10 each). CpoS-FLAG constructs were precipitated from lysates at 26 hpi. Eluates were analyzed by western blot analysis (* indicates visible Co-IP of IPAM(L2)-MYC). Note, the bands of CpoS(L2-CC1*) and CpoS(L2-CC1/2*) (stained with anti-FLAG) overlap with the band of the co-eluted IgG light chain, and the band of CpoS(Cm) is weak. **(B)** Localization of IPAM in infected HeLa cells (MOI5, 22 hpi, scale=20 µm). **(C)** *Chlamydia*’s ability to dampen cell death and IFN responses does not depend on IPAM. Luciferase activity in A2EN-ISRE reporter cells (MOI5, 14 hpi) and cell death in A2EN and HeLa cells (MOI5, 24 hpi) (mean±SD, n=3-7, one-way ANOVA: indicated are significant differences compared to CTL2).

Comparing the differential abilities of CpoS(L2-CC1*) and CpoS(L2-CC2*) to restore proper IPAM localization (Fig 4B) with their differing abilities to block host cell death and IFN responses (Fig 2D-F), it seems unlikely that IPAM (mis)localization and its consequences on the MT cytoskeleton are directly linked to the induction of host cell defense responses. Indeed, neither the IPAM-MYC-expressing strain, which overexpresses IPAM, nor a strain deficient for IPAM (CTL2-*ipaM*::*bla*) (Fig S2A), both of which express CpoS, differed from the wild-type CTL2 strain with respect to host cell killing or induction of IFN responses (Fig 4C). In contrast, a CpoS/IPAM double-deficient strain (CTL2-*ipaM*::*bla-cpoS::cat*) (Fig S2B) behaved like a CpoS single-deficient strain and hence induced premature cell death and an exaggerated IFN production (Fig 4C). Interestingly, the more pronounced formation of MT cages observed during infection with CTL2-*cpoS*::*cat* (Fig 3G), was independent of IPAM and its mislocalization, as it could also be observed in cells infected with the CpoS/IPAM double-deficient strain, but for instance not in cells infected with the IPAM-MYC expressing strain, which overexpresses IPAM (Fig S2C).

Overall, these observations show that the role of CpoS in blocking cell-autonomous defenses is not linked to its role in maintaining inclusion microdomains.

### CpoS mediates the recruitment of Rab GTPases and modulates membrane trafficking

Numerous members of the Rab GTPase family, which influence membrane trafficking and organelle identity (Hutagalung et al., 2011), are recruited to the CTL2 inclusion (Rzomp et al., 2003). We and others have previously shown that CpoS(L2) and CpoS(D) interact with multiple Rab GTPases (Mirrashidi et al., 2015, Rzomp et al., 2006, Sixt et al., 2017). Of note, the recruitment of Rab1A to the CTL2 inclusion depends on CpoS (Sixt et al., 2017), and other Rab proteins might be recruited in a CpoS-dependent fashion as well (Faris et al., 2019).

To confirm that CpoS is required for the recruitment of a diverse set of Rab GTPases, we monitored the localization of transiently expressed EGFP-Rab fusion proteins in infected HeLa cells. At 24 hpi, Rab1A, 4A, and 35 were prominently enriched at the periphery of CTL2 inclusions (Fig 5A and S3A). In contrast, consistent with a previous report (Rzomp et al., 2003), Rab5A was not recruited to inclusions, although Rab5A-containing vesicles could be seen in the vicinity of some inclusions (Fig S3A). Significantly, the broad recruitment of Rab1A, 4A, and 35 was absent in cells infected with CTL2-*cpoS*::*cat*, but was restored in cells infected with the complemented strain expressing CpoS(L2) (Fig 5A and S3A).

**Figure 5.**
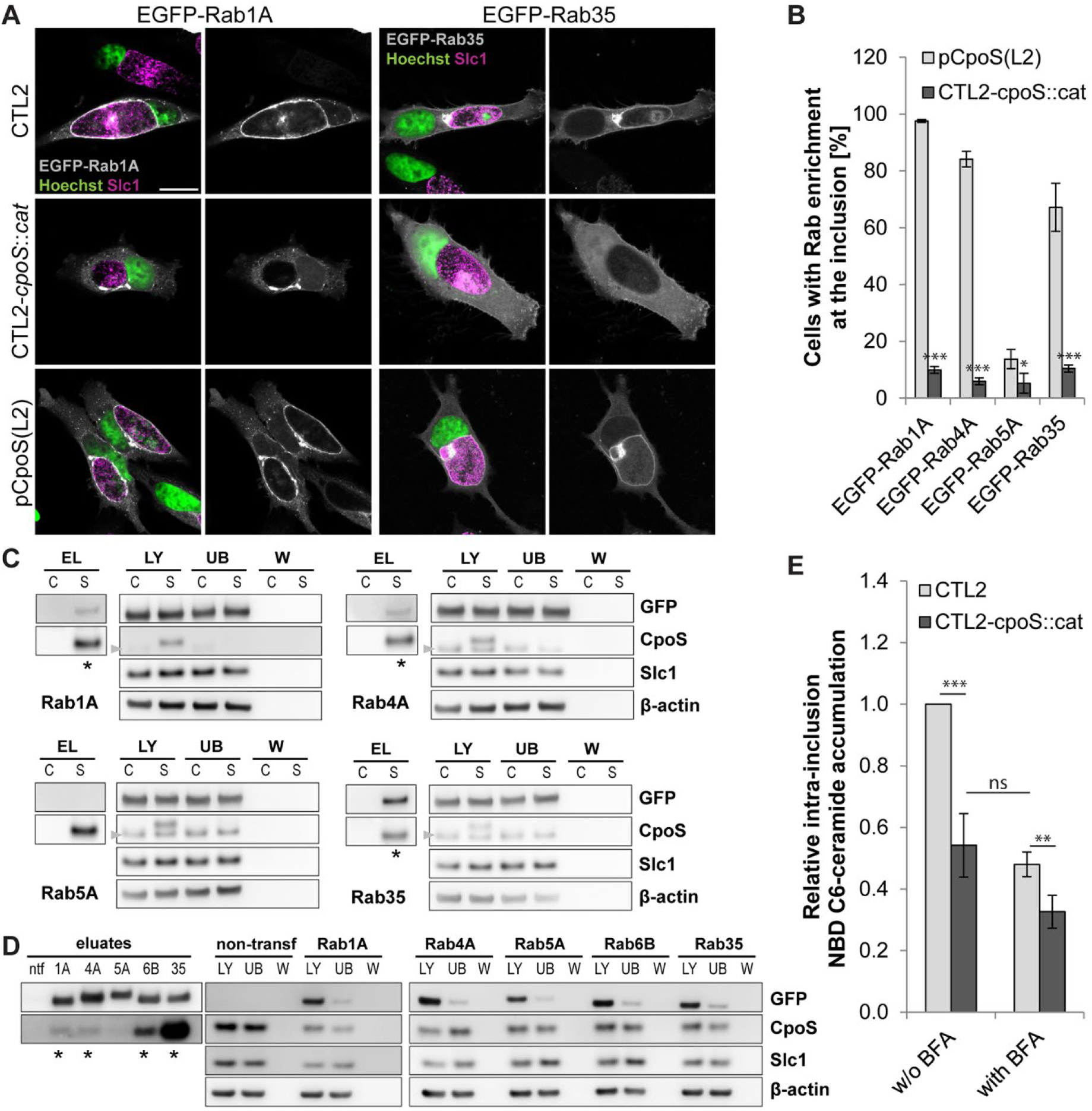
CpoS mediates the recruitment of Rab GTPases and modulates membrane trafficking. **(A)** Localization of EGFP-tagged Rab1A and 35 in infected HeLa cells (MOI5, 24 hpi, confocal, scale=20 µm). **(B)** Percentage of infected HeLa cells with enrichment of EGFP-Rab proteins at the inclusion (MOI5, 24 hpi). BFA (3 µg/ml) added at 18 hpi (mean±SD, n=3 (at least 250 cells per condition), student t test). **(C-D)** Co-IP confirms interaction between CpoS and Rab GTPases. HeLa cells were infected (MOI20; CTL2 (=C(ontrol)) or CTL2-*cpoS*::*cat*/pCpoS(L2)-FLAG (=S(ample)) in (C); CTL2-*cpoS*::*cat*/pCpoS(L2)-FLAG in (D)) and transfected with EGFP-Rab expression plasmids (ntf, non-transfected). CpoS-FLAG (C) or EGFP-Rab proteins (D) were precipitated from lysates at 26 hpi. Samples were analyzed by western blot analysis (LY, lysate; UB, unbound; W, wash; EL, eluate; * visible Co-IP). Note that the CpoS-specific antibody used in (C) detects both endogenous and FLAG-tagged CpoS, yet also a non-specific band (also seen in uninfected cells (not shown)) that overlaps with the band of endogenous CpoS (arrowheads). **(E)** Reduced ceramide acquisition by CpoS-deficient strain. At 14 hpi, infected HeLa cells (MOI10) were treated with NBD C6-ceramide, in presence or absence of BFA (3 µg/ml). At 21 hpi, the average intra-inclusion ceramide fluorescence intensity was determined (mean±SD, n=3, one-way ANOVA).

We frequently observed an additional accumulation of Rab1A-positive membranes adjacent to discrete parts of the inclusion, a phenomenon also seen in cells infected with CTL2-*cpoS*::*cat* (Fig 5A). This can be explained by the natural enrichment of Rab1A at the Golgi apparatus, an organelle known to fragment in *Chlamydia*-infected cells, generating Golgi mini-stacks that accumulate at the inclusion periphery (Heuer et al., 2009). Indeed, treatment with brefeldin A (BFA), a drug that disrupts normal Golgi morphology (Donaldson et al., 1992, Helms et al., 1992), dispersed these structures, while leaving Rab1A recruitment to CTL2 inclusions unaffected (Fig S3B). To distinguish between Golgi and inclusion membrane association, we quantified the percentage of infected cells displaying recruitment of selected Rab GTPases to inclusions in cells treated with BFA (Fig 5B). Moreover, to complement the microscopic observations, we conducted Co-IPs. EGFP-tagged Rab1A, 4A, and 35, but not 5A, could be co-precipitated with CpoS(L2)-FLAG from lysates of infected HeLa cells using anti-FLAG beads (Fig 5C). Conversely, CpoS(L2)-FLAG could be co-precipitated with EGFP-tagged Rab1A, 4A, and 35 using beads coated with anti-GFP-antibodies (Fig 5D).

When we expanded our microscopic studies to additional Rab proteins, we found that recruitment of Rab4B and 14 also depended on CpoS (Fig S3C). In contrast, Rab18 was recruited neither to CpoS-containing nor to CpoS-deficient inclusions, while recruitment of Rab8A and 10 was seen in some cells, but occurred independently of CpoS (Fig S3C). Moreover, the accumulation of Rab6A and 6B in the vicinity of inclusions was occasionally observed, yet appeared to be primarily a consequence of the recruitment of Golgi fragments to inclusions, and no clear difference was seen in the presence or absence of CpoS (Fig S3C). This latter finding was unexpected, as our Co-IP experiments detected an interaction between CpoS and Rab6B (Fig 5D).

The recruitment of Rab proteins has been proposed to reflect the ability of the bacteria to capture host vesicles and hence to acquire lipids (Damiani et al., 2014, Faris et al., 2019). In support of this concept, we observed that the trafficking of NBD C6-ceramide (or derived labeled lipids) to inclusions was reduced in the absence of CpoS (Fig 5E). The level of reduction was comparable to that seen when vesicular transport was disturbed by addition of BFA (Fig 5E).

Taken together, these data confirm that CpoS recruits a diverse set of Rab GTPases to the *C. trachomatis* inclusion and that the absence of CpoS compromises the capacity of *C. trachomatis* to subvert the membrane trafficking machinery of its host cell.

### Rab35 participates in the CpoS-mediated blockade of the IFN response

To explore a potential link between Rab recruitment and the subversion of host cell defenses by CpoS, we determined whether CpoS orthologs and variants would restore interactions with Rab proteins.

When expressed in CTL2-*cpoS*::*cat*, all the tested CpoS orthologs restored the recruitment of Rab1A, 4A, and 35 in infected HeLa cells (Fig S4A). Rab recruitment required interactions mediated by CC1 (or residues therein), as CpoS(L2-CC1*) and CpoS(L2-CC1/2*) failed to recruit any of the three Rab GTPases, contrasting with CpoS(L2-CC2*), which did restore recruitment of Rab1A, 4A, and 35 (Fig 6A-B and S4B). CpoS(L2Δ60), which lacks the far C-terminus of CpoS, also lost its ability to recruit Rab1A and 4A, while it was still capable of recruiting Rab35 (Fig 6A-B). Co-IP experiments supported our microscopic observations in respect to the restoration of Rab recruitment by the distinct orthologs and variants of CpoS (Fig 6C). Moreover, consistent with a role for CpoS-Rab interactions in ceramide acquisition, CpoS(L2) could restore normal levels of ceramide acquisition when expressed in CTL2-*cpoS*::*cat*, while the modified variants of CpoS(L2) failed to do so, with the exception of CpoS(L2-CC2*) (Fig 6D), the only variant tested that restored interactions with a broad range of Rab proteins (Fig 6A-B and S4B).

**Figure 6.**
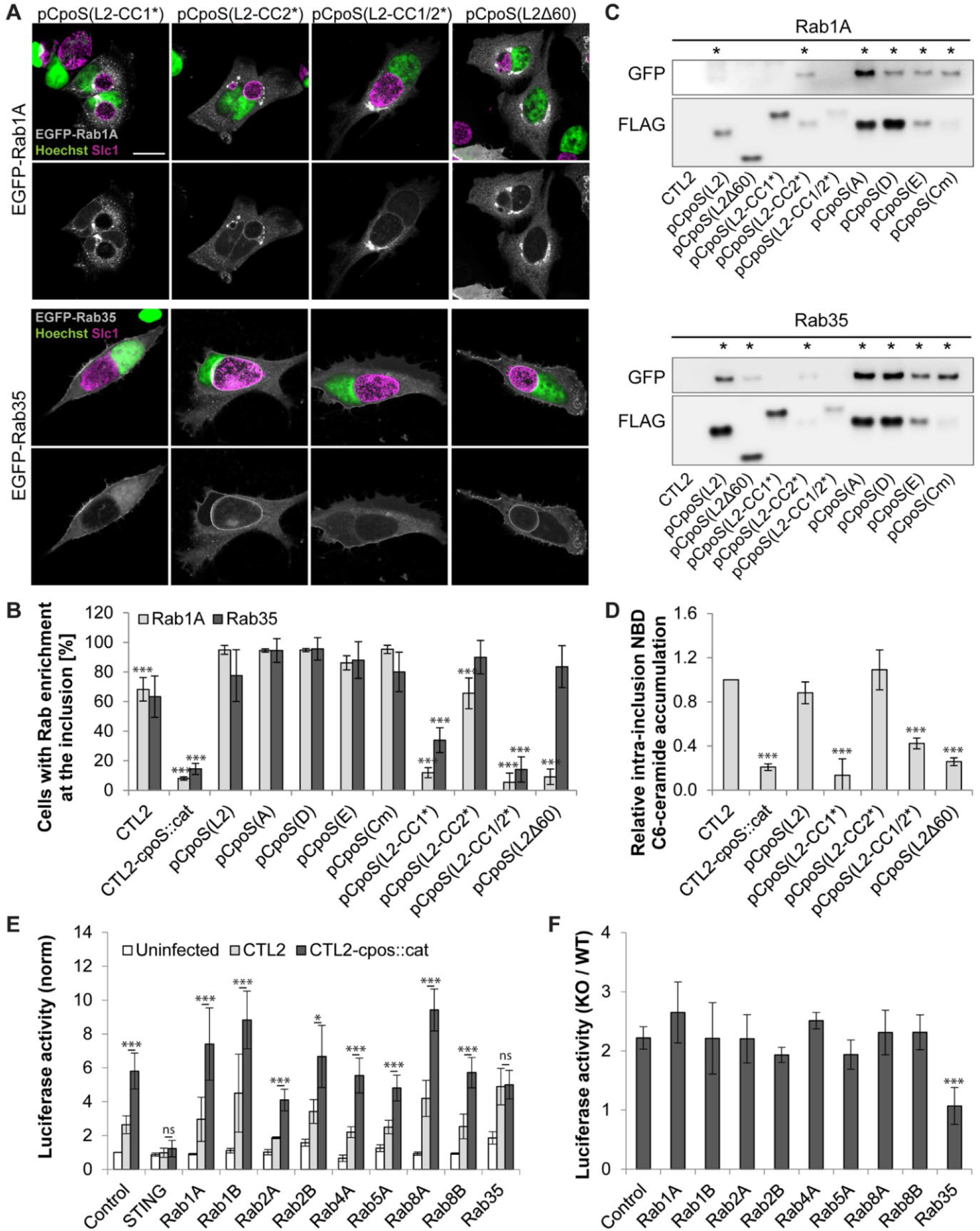
Rab35 participates in the CpoS-mediated blockade of the IFN response. **(A)** Localization of EGFP-tagged Rab1A and 35 in infected HeLa cells (MOI5, 24 hpi, confocal, scale=20 µm). **(B)** Enrichment of EGFP-tagged Rab1A and 35 at inclusions (MOI5, 24 hpi). BFA (3 µg/ml) was added at 18 hpi (mean±SD, n=3 (at least 75 cells per condition), one-way ANOVA: significant differences compared to pCpoS(L2)). **(C)** Co-IP reveals the differential ability of CpoS variants to interact with EGFP-Rab proteins. CpoS-FLAG variants were precipitated from lysates of infected HeLa cells (MOI20, 26 hpi). Eluates were analyzed by western blot analysis (* visible Co-IP). **(D)** Differential ability of CpoS variants to restore ceramide acquisition. At 14 hpi, infected HeLa cells (MOI10) were treated with NBD C6-ceramide. At 21 hpi, the average intra-inclusion ceramide fluorescence intensity was determined (mean±SD, n=3, one-way ANOVA: significant differences compared to CTL2). **(E-F)** Effect of siRNA-mediated Rab depletion on the inhibition of the IFN response in infected A2EN-ISRE reporter cells (MOI10, 14 hpi). Displayed are normalized luciferase activities (E), as well as the corresponding ratio of activities detected in CTL2-*cpoS*::*cat* and CTL2 (F) [mean±SD, n=3-7, one-way ANOVA: significant differences compared to control (F) or as indicated (E)].

When compared to the differential abilities of CpoS(L2-CC1*) and CpoS(L2-CC2*) to block the host cell-autonomous defense, these findings indicated a correlation between the suppression of the IFN response by CpoS and its ability to recruit Rab proteins. Indeed, only those variants that could mediate Rab interactions were capable of dampening the IFN response (compare Fig 1F and 2E-F with Fig 6B). To elucidate the potential role of Rab proteins in the regulation of the IFN response, we evaluated the effect of siRNA-mediated depletion of individual Rab proteins on IFN-driven luciferase production in the A2EN reporter cells. We included siRNAs targeting the main Rab proteins considered earlier in this study, as well as a few others. As expected, depletion of STING, included as a control, completely blocked luciferase production in response to either CTL2 or CTL2-*cpoS*::*cat* (Fig 6E). Depletion of individual Rab proteins had variable effects. For instance, depletion of Rab1B or 8A enhanced, but that of Rab2A reduced, IFN-driven luciferase production during infection (Fig 6E). Of note, the depletion of individual Rab proteins did not affect the increase in IFN signaling observed during infection with CTL2-*cpoS*::*cat* compared to CTL2 (Fig 6F). A notable exception was the depletion of Rab35, which compromised the ability of CTL2 to dampen luciferase production to a level that resembled those induced by the CpoS-deficient strain CTL2-*cpoS*::*cat* (Fig 6E-F and S4C). Hence, with respect to the IFN response quantified by the luciferase reporter, Rab35 depletion in host cells phenocopied the effects of CpoS deficiency in *C. trachomatis*.

Altogether, our findings indicate that CC1 and the far C-terminus of CpoS mediate interactions with Rab GTPases and that Rab35 plays an important role in the CpoS-mediated dampening of the IFN response during *Chlamydia* infection.

## DISCUSSION

In this study, we characterized the interaction partners and biological functions of orthologs and modified variants of CpoS with the intention of determining the mechanism by which CpoS subverts the host cellular defense. We made two key discoveries (Fig 7): (a) CpoS is essential for the proper formation and/or maintenance of inclusion microdomains. Its absence severely disrupts the localization of multiple inclusion-associated proteins, thus significantly altering the microenvironment of *Chlamydia*’s parasitophorous vacuole (Fig 3). (b) The role of CpoS in suppressing the host type I IFN response is independent of its function in inclusion microdomains (Fig 2 and 4). Instead, it is linked to the ability of CpoS to modulate membrane trafficking via its interactions with Rab proteins, such as Rab35 (Fig 2 and 6).

**Figure 7.**
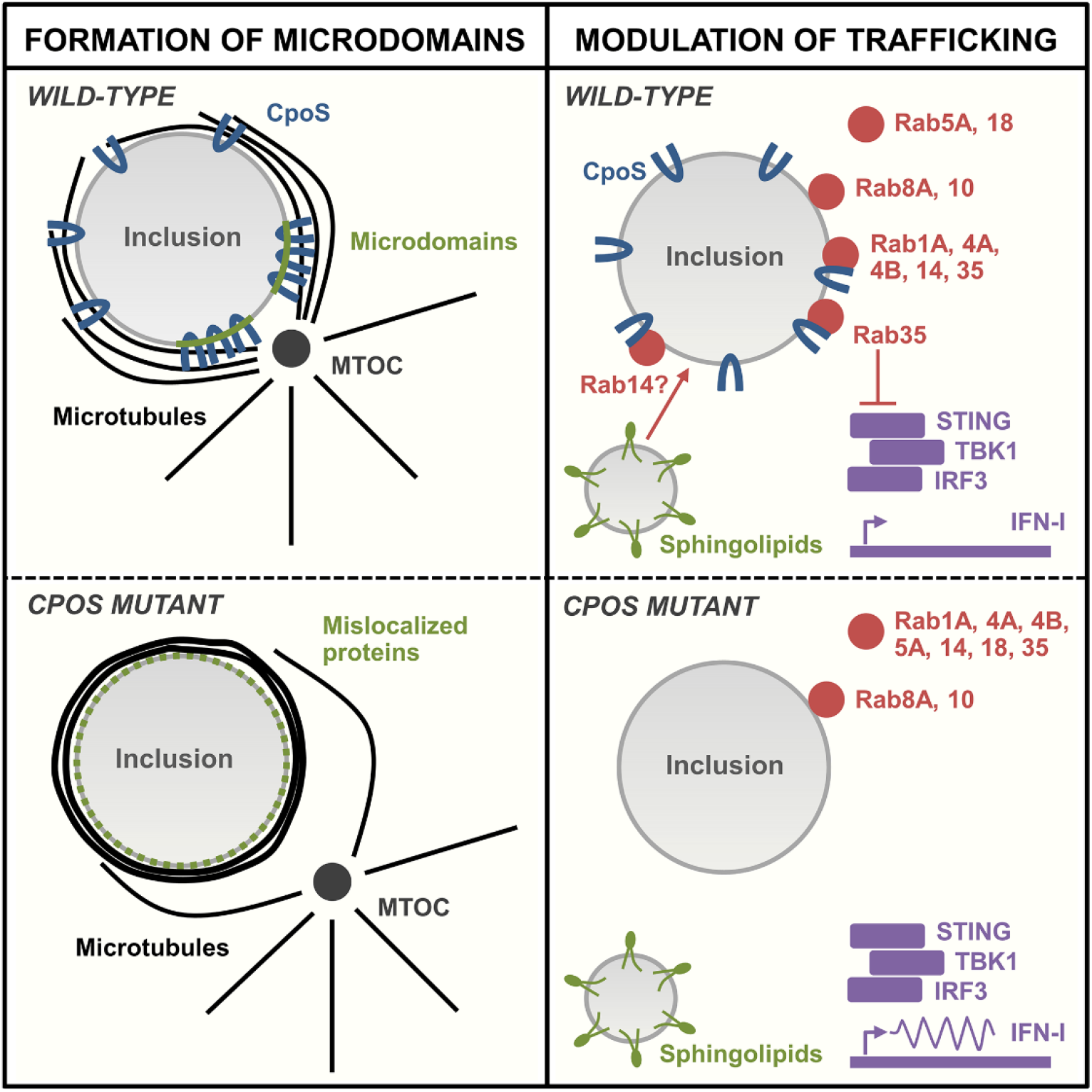
Graphical summary of the role of CpoS in inclusion microdomains and modulation of membrane trafficking and resulting implications.

Many biological membranes form microdomains, i.e. patches of distinct composition that serve special functions in signaling and trafficking (Bagam et al., 2017). In mammalian cells, these domains can also have roles in mediating host-pathogen interactions. For example, pathogens often exploit lipid rafts, cholesterol-rich microdomains in the plasma membrane of their host cells, as entry points during invasion and to hijack host cell signaling pathways (Samanta et al., 2017). In addition, microdomains have been observed in the vacuolar membranes of diverse intracellular bacterial pathogens (Howe et al., 2006, Knodler et al., 2003, Samanta et al., 2017).

In *C. trachomatis* inclusions, microdomains have been described as discrete cholesterol-rich patches that co-localize with activated Src kinases and multiple Incs, including CTL0223 (CT850), CTL0356 (CT101/MrcA), CTL0475 (CT222), CTL0476 (CT223/IPAM), CTL0477 (CT224), CTL0480 (CT228), CTL0484 (CT232/IncB), CTL0485 (CT233/IncC), and CTL0540 (CT288) (Lutter et al., 2013, Mital et al., 2010, Weber et al., 2015). Our finding that CpoS regulates the formation and/or maintenance of these microdomains was unexpected, as CpoS had not previously been reported to associate with these structures. Moreover, Src-enriched microdomains have been observed in inclusions inhabited by *C. pneumoniae*, but not in inclusions of other *Chlamydia* species such as *C. caviae* and *C. muridarum* (Mital et al., 2010), while, among these species, a CpoS ortholog can be found only in *C. muridarum*. Hence, CpoS cannot be the sole factor that determines the generation of inclusion membrane microdomains across *Chlamydia* spp. Nevertheless, the fact that CpoS(Cm) restored localization of IPAM to microdomains when expressed in the *cpoS* mutant (Fig 4B), indicates that a conserved CpoS function can control the occurrence of microdomains.

Inclusion microdomains have been proposed to act as hubs that regulate the trafficking of inclusions to, and their stable association with, centrosomes (Mital et al., 2011, Mital et al., 2015), the MT architecture (Dumoux et al., 2015), and bacterial exit by extrusion (Lutter et al., 2013, Nguyen et al., 2018). Many more functions likely remain to be discovered. Hence, the mislocalization of multiple host and bacterial microdomain-associated proteins in cells infected with *cpoS* mutants is expected to compromise numerous interactions between the host and inclusions. Consistent with this prediction, we observed that the MT cage surrounding CpoS-deficient inclusions is altered (Fig 3G). Of note, these specific changes were independent of (the mislocalization of) IPAM (Fig S2C), an Inc previously reported to act on MTs (Dumoux et al., 2015), suggesting a role for additional microdomain-associated proteins in the modulation of the host cytoskeleton. Because we have not characterized the subcellular localization of each of the previously reported microdomain-associated Incs, we cannot exclude that some of the Incs accumulate in altered types of microdomains, nor can we exclude that some of the Incs retain some functionality in the absence of microdomains.

While CpoS deficiency caused a striking disruption of inclusion microdomains, our studies suggested that the CpoS-mediated suppression of the host type I IFN response was independent of its role in microdomains, but instead dependent on its interactions with Rab GTPases via its CC1 motif. Indeed, expression of CpoS(L2-CC1*) restored interactions with IPAM and the localization of IPAM to microdomains (Fig 4A-B), but failed to restore interactions with Rab proteins (Fig 6A-C) or the blockade of the IFN response (Fig 2E-F). The opposite phenotypes were observed for CpoS(L2-CC2*) (Fig 2E-F, 4A-B and 6A-C).

The selective recruitment of Rab proteins to the *Chlamydia* inclusion is a well-studied phenomenon (Rzomp et al., 2003), but a role for CpoS in this process has only recently been confirmed genetically. We previously described that Rab1A recruitment depends on CpoS (Sixt et al., 2017), and now expand this finding to Rab4A, 4B, 14 and 35 (Fig 5A-B and S3), confirming CpoS as a broad recruiter of Rab proteins (Faris et al., 2019). We also found that CpoS-Rab interactions require an intact CC1 region and that the interactions with some Rab proteins, such as Rab1A and 4A, but not Rab35, additionally depend on the far C-terminus of CpoS (Fig 6A-C). These observations are consistent with recent reports regarding the implication of residues 120 (located in CC1) and residues 197-215 (located at the far C-terminus) in CpoS-Rab interactions (Faris et al., 2019, Rzomp et al., 2006). However, although Rab6 recruitment to *C. trachomatis* inclusions has been reported (Faris et al., 2019, Moorhead et al., 2007, Rzomp et al., 2003), and our Co-IP studies suggested an interaction between CpoS and Rab6B (Fig 5D), we could not validate such an interaction by microscopy (Fig S3C). Moreover, at difference with a prior report (Faris et al., 2019), we did not find a clear CpoS-dependence for the recruitment of Rab8A and 10 (Fig S3C), suggesting that other factors may be required to relocalize these proteins to inclusions.

Modulation of Rab localization or activity allows intracellular bacteria to control the trafficking of their vacuole and its interactions with host cellular compartments (Brumell et al., 2007). Endocytic Rab proteins are excluded from the *Chlamydia* inclusion presumably to avoid lysosomal targeting of the bacteria (Rzomp et al., 2003). Others may be recruited to enable the capture of vesicles that contain lipids important for bacterial replication (Damiani et al., 2014). Indeed, depletion or inhibition of certain Rab proteins, including Rab14, which we found to be recruited by CpoS (Fig S3C), decreases ceramide (sphingolipid) transport to inclusions and the generation of infectious EBs (Capmany et al., 2010, Gambarte Tudela et al., 2015, Rejman Lipinski et al., 2009). Consistently, we found ceramide acquisition by inclusions reduced in absence of CpoS (Fig 5E), confirming the functional significance of CpoS-Rab interactions.

While we found that only those variants of CpoS that recruited Rab proteins were capable of dampening STING activation and ISRE-driven gene expression (Fig 2E-F and 6A-C), we do not know if CpoS directly interferes with the trafficking of STING, which is essential for the activation and termination of its signaling function (Dobbs et al., 2015, Gonugunta et al., 2017). Alternatively, a loss of the capacity to modulate membrane trafficking by CpoS may affect the composition and stability of the inclusion, resulting in enhanced leakage of bacterial components from inclusions, thus causing enhanced STING activation. Our work also identified a specific role for Rab35, a protein engaged in endocytic recycling (Klinkert et al., 2016), in the CpoS-mediated inhibition of the IFN response (Fig 6E-F). This is interesting, because various intracellular bacteria modulate Rab35 activity or localization (Bakowski et al., 2010; Dikshit et al., 2015; Furniss et al., 2016; Huang et al., 2010; Mukherjee et al., 2011), implying that the suppression of the IFN response through exploitation of Rab35 function could be a more widespread pathogenic strategy. Of note, our observation that CpoS(L2Δ60) could restore Rab35 recruitment (Fig 6A-B), but not the inhibition of the IFN response (Fig 2E-F), suggests that Rab35 needs to act together with additional factors recruited by CpoS, potentially other Rab proteins, to dampen the defense response.

Interestingly, we did not observe strong evidence for a connection between CpoS-Rab interactions and the inhibition of the premature death that occurs in cells infected with *cpoS* mutants. Indeed, both CpoS(L2)-CC1* and CpoS(L2)-CC2* conferred intermediate levels of protection against cell killing (Fig 2D), suggesting that functions mediated by both CC motifs are required for full cytoprotection. This agrees with our previous finding that the cell death response does not require a functional IFN response (Sixt et al., 2017), meaning that CpoS does not block cell death solely by interfering with STING activation. Restoring Rab recruitment and inhibition of the IFN response, by expression of CpoS(L2)-CC2*, only marginally improved the formation of infectious EBs (Fig 2G), consistent with the premise that the disruption of EB generation in absence of CpoS is primarily caused by the premature loss of the host cell. However, it is plausible that the inhibition of the IFN response has implications for the pathogenicity of *in vivo* infections, in which the subversion of non-cell-autonomous immune responses involving specialized immune cells should increase the virulence of *Chlamydia*.

Altogether, this work highlights the importance of Inc-Inc associations in shaping the microenvironment of *Chlamydia*’s parasitophorous vacuole, and uncovers the importance of CpoS-Rab interactions for the evasion of the host cellular defense by *C. trachomatis*. Moreover, these mechanistic insights can support future efforts to explore and exploit CpoS as a potential drug target.

## MATERIALS AND METHODS

### *In silico* analysis of CpoS

Orthologs of CpoS(L2) were identified using blastp (Altschul et al., 1990). Pairwise sequence identities were determined using EMBOSS Needle at EMBL-EBI (Madeira et al., 2019), while the multiple sequence alignment was made with Clustal Omega (Madeira et al., 2019) using the Boxshade 3.21 tool at Expasy (Duvaud et al., 2021) for formatting. Protein topology was predicted using the Phobius tool (Kall et al., 2004) via InterProScan 5 (Jones et al., 2014). Coiled-coil motifs were predicted using Paircoil2 (Berger et al., 1995). Disruptive effects of amino acid substitutions on coiled-coil motifs were predicted using both Paircoil2 (Berger et al., 1995) and Coils (Lupas et al., 1991).

### Mammalian cell culture

HeLa (ATCC CCL-2) and Vero (ATCC CCL-81) cells, as well as immortalized *Gt* MLFs expressing wild-type STING-HA (Barker et al., 2013), were grown in Dulbecco’s Modified Eagle’s Medium (DMEM; Gibco) supplemented with 10% heat-inactivated fetal bovine serum (FBS; Gibco). In selected experiments, i.e. experiments including an analysis of cell death or the imaging of live cells, a DMEM medium devoid of phenol red was used to avoid interference of phenol red with measurements of absorbance or fluorescence. A2EN (Buckner et al., 2011) and A2EN-ISRE reporter cells (Sixt et al., 2017) were cultivated in Keratinocyte-SFM (Gibco) supplemented with 10% heat-inactivated FBS. Cultures were maintained in a humidified incubator (37°C, 5% CO_2_) and were routinely tested for *Mycoplasma* contamination using a PCR assay described elsewhere (van Kuppeveld et al., 1992) or by using a commercial PCR-based *Mycoplasma* detection assay (VenorGeM, Minerva).

### Transient expression in human cells

Plasmids enabling transient expression of EGFP-tagged human Rab proteins were kindly provided by Marci Scidmore (formerly at Cornell University). The generation of these plasmids, which are derivatives of pEGFPC1 and pEGFPC2 (Clontech), was described in previous publications (Rzomp et al., 2003; Huang et al., 2010). To confirm the identity of the cloned Rab genes, plasmid inserts were sequenced (Eurofins or GATC sequencing services) using primer pEGFPC1for (Table S2) and the sequences were compared with respective mRNA sequences available at NCBI gene. Plasmid pcDNA3.1-Cyto-GFP1-10, enabling transient expression of cytosolic GFP1-10 for the detection of GFP11-tagged proteins, was kindly provided by Kevin Hybiske (University of Washington). In experiments including infection with *Chlamydia*, cells were transfected with these plasmids either 2-4 hours before infection or at 1 hpi using jetPRIME transfection reagent (Polyplus), with a medium exchange after 2-3 hours, or lipofectamine 2000 (Invitrogen), without medium exchange.

### *Chlamydia* strains and infection procedure

Experiments were carried out with *C. trachomatis* strain L2/434/Bu (CTL2, ATCC VR-902B) and the derivatives generated in this study (Table S1). For the generation of bacterial stocks used for infection experiments, bacteria were harvested from infected Vero cells by H_2_O-mediated lysis, as described previously (Sixt et al., 2017), and stored in 1xSPG (sucrose-phosphate-glutamate) buffer (75 g/l sucrose, 0.5 g/l KH_2_PO_4_, 1.2 g/l Na_2_HPO_4_, 0.72 g/l glutamic acid, pH 7.5) at −80°C. Bacteria were shown to be free of *Mycoplasma* contamination by PCR, as described above for cell lines. To determine the number of infectious bacteria (i.e. the inclusion-forming units (IFUs)) contained in the bacterial stocks, Vero cell monolayers were infected with serial dilutions of the bacteria. At 28 hpi, cells were fixed with 4% formaldehyde and inclusions were stained with antibodies targeting the chlamydial protein Slc1 (procedure described below). Inclusions were counted using a Cellomics ArrayScan VTI HCS automated imaging system (Thermo-Fisher-Scientific). To conduct infections, cells were seeded in multi-well plates, followed by addition of bacteria (number of IFU/cell, also known as multiplicity of infection (MOI), as specified), centrifugation (1500 x g, 30 min, 23°C), and incubation (37°C, 5%CO_2_) for the indicated periods of time.

### Gene disruption in *Chlamydia*

CTL0476 (*ipaM*) was disrupted in CTL2 using the TargeTron approach (Johnson et al., 2013), similarly as recently described for disruption of CTL0481 (*cpoS*) (Sixt et al., 2017). The primers IBS-CTL0476, EBS1d-CTL0476, and EBS2-CTL0476 (*Table S2*), as well as EBS universal (Sigma-Aldrich), were used to retarget vector pDFTT3 (Johnson et al., 2013) for this purpose. To disrupt CTL0481 with an intron carrying a chloramphenicol resistance marker, vector pDFTT3-CTL0481 (Sixt et al., 2017) was modified in a two-step process. First, vector pDFTT3-CTL0481 was PCR-amplified (QuikChange II XL Kit, Agilent) using primers pDFTT3-rvcat_F and pDFTT3-rvcat_R (Table S2), digested with ApaI (NEB), and circularized by ligation (T4 DNA ligase, NEB) to remove the *cat* gene from the vector backbone. Second, this modified vector was further PCR-amplified with primers pDFTT3-rvbla_F and pDFTT3-rvbla_R (Table S2), digested with NdeI and SphI (NEB), and ligated to a NdeI/SphI-digested gene block (containing the *cat* gene; IDT) to replace the *bla* gene with the *cat* gene in the intron. CaCl_2_-mediated transformation of *Chlamydia* (Wang et al., 2011) was conducted as recently described (Sixt et al., 2017). Transformants were selected in the presence of 1 U/ml penicillin G (Sigma-Aldrich) or 0.5 µg/ml chloramphenicol (Sigma-Aldrich), first added at 12 hpi. Transformants were plaque-purified in presence of 5 U/ml penicillin G or 1 µg/ml chloramphenicol. Intron insertion at correct target sites was verified by PCR (using primer pairs CTL0476_seqF/R or CTL0481_seqF/R, Table S2) and by sequencing of the resulting PCR products (Eton Bioscience).

### Expression of CpoS orthologs and variants

To enable expression of CpoS orthologs and variants with C-terminal FLAG tag, a DNA fragment encoding a FLAG tag and a stop codon was introduced between the KpnI and SalI restriction sites of the *E. coli*-*Chlamydia* shuttle vector p2TK2-SW2 (Agaisse et al., 2013), generating vector p2TK2-SW2-FLAG. DNA fragments coding for CpoS orthologs and variants (without stop codon) and their promoter regions (210 bp sequence upstream of the start codon) were PCR-amplified from genomic DNA or were obtained as synthetic gene blocks (IDT) and were subsequently inserted between the AgeI and KpnI restriction sites of vector p2TK2-SW2-FLAG. The following primers were used for PCR amplification (Table S2): CTL0481 (*C. trachomatis* L2/434/Bu, CTL0481_F and CTL0481_R), CTL0481Δ60 (*C. trachomatis* L2/434/Bu, CTL0481_F and CTL0481Δ60_R), CTA0251 (*C. trachomatis* A/HAR-13, CTL0481_F and CTA0251_R), CT229 (*C. trachomatis* D/UW-3/CX, CTL0481_F and CTL0481_R), BOUR00241 (*C. trachomatis* E/Bour, CTL0481_F and CTL0481_R), and TC0500 (*C. muridarum* Nigg, TC0500_F and TC0500_R). Vectors were then transformed into strain CTL2-*cpoS*::*cat* (as described above). The presence of the correct constructs in plaque-purified transformed bacteria was confirmed by PCR (using primers p2TK2-SW2_seqF and p2TK2-SW2_seqR, Table S2) and sequencing of resulting PCR products (Eton Bioscience).

### Heterologous gene expression in *Chlamydia*

To generate vector p2TK2-SW2-IPAM-MYC, enabling expression of MYC-tagged-IPAM, CTL0476 (*ipaM*) and its promoter region (210 bp sequence upstream of the start codon) were PCR-amplified from genomic DNA of *C. trachomatis* L2/434/Bu with a primer pair that introduces a C-terminal MYC-tag (CTL0476-MYC_F and CTL0476-MYC_R, Table S2). The PCR product was digested with SgrAI and NheI and subsequently inserted between the AgeI and NheI restriction sites of vector p2TK2-SW2 (Agaisse et al., 2013). Vector p2TK2-SW2-mCherry (Agaisse et al., 2013), enabling constitutive expression of mCherry, was a kind gift from Isabelle Derré (University of Virginia). Vector pTL2-tetO-IncB-GFP11x7-flag, enabling inducible expression of IncB (CT232) tagged at its C-terminus with seven copies of GFP11 (RDHMVLHEYVNAAGIT) and one copy of FLAG (DYKDDDDK), was a kind gift from Kevin Hybiske (University of Washington). The vectors were transformed into strains CTL2 and/or CTL2-*cpoS*::*cat* (as described above). In experiments using IncB-GFP11x7-FLAG-expressing strains, expression was induced by addition of 4 ng/ml anhydrotetracycline (Clontech) at 4 hpi.

### Quantification of cell death and IFN response

LDH activity in culture supernatants, an indicator for host cell lysis, was quantified using the photometric *in vitro* cytotoxicity kit (Sigma-Aldrich), according to the manufacturer’s instructions. Activity detected in cell-free medium (blank) was subtracted and values were normalized to activity detected in a total cell lysate to calculate the percentage of dead cells. The induction of type I IFN-dependent genes was assessed using an A2EN-ISRE-luciferase reporter cell line (Sixt et al., 2017). Cells were lysed by addition of a 1:1 mixture of Hanks’ Balanced Salt Solution (HBSS; Gibco) and Britelite plus reagent (PerkinElmer), before measurement of luminescence. Luminescence values were normalized to the mean luminescence detected in uninfected (non-siRNA-treated) wells. All absorbance and luminescence measurements for the assays described above were conducted using a plate reader, such as an EnSpire 2300 (PerkinElmer), SpectraMax i3 (Molecular Devices) or Infinite 200 (Tecan). The same instrument was used for all replicates of the same experiment.

### Determination of infectious progeny

To quantify the formation of infectious progeny, 96-well plates with confluent monolayers of HeLa cells were infected with a low dose of the indicated *C. trachomatis* strains (< 50% infected cells). At 36 hpi, cell lysates were prepared by H_2_O-based cell lysis. Infectious particles in the initial inoculum (input) and collected cell lysates (output) were quantified by infecting confluent monolayers of Vero cells with serial dilutions, followed by the fluorescence microscopic determination of inclusion numbers at 28 hpi (as described above for the titering of bacterial stocks). From the IFUs detected in the input and the output, the number of infectious particles formed per infected cell could be determined for each strain, and was then normalized to the IFU production observed for CTL2.

### Ceramide acquisition

To measure the intra-inclusion accumulation of ceramide (and/or ceramide-derived lipids), cells were seeded in 96-well plates and infected with the indicated strains (for the experiment shown in Fig 5E, mCherry-expressing derivatives of the indicated strains were used). At 14 hpi, cells were incubated for 15 min at 4°C, rinsed once with cold HBSS (Gibco), and then incubated for 30 min at 4°C in HBSS containing 5 µM NBD C6-ceramide complexed to BSA (Thermo-Fisher-Scientific). Subsequently, cells were rinsed twice with pre-warmed (room temperature) growth medium and incubated for another 6 h in growth medium at 37°C, 5% CO_2_. In some wells, BFA (3 µg/ml) was added to the culture medium at 12 hpi and was also present during the incubation with ceramide, but not thereafter. At the end of the 6-hour incubation period, cells were stained for 10 min in growth medium containing 2 µg/ml Hoechst 33342, washed twice with growth medium, and imaged live with an automated fluorescence microscope (ImageXpress Micro XL, Molecular Devices) at about 21 hpi. Images were analyzed with the software MetaXpress (Molecular Devices). Inclusions were detected based on the fluorescence of mCherry (in experiments using mCherry-expressing strains) or fluorescence of NBD (in all other experiments). The average intra-inclusion NBD fluorescence intensity was determined and normalized to intra-inclusion NBD fluorescence observed during infection with CTL2.

### siRNA-mediated gene silencing

Cells were transfected with siRNAs (Dharmacon siGENOME Human SMART pools, i.e. mix of four gene-specific siRNAs, Table S3) according to the manufacturer’s guidelines, using DharmaFECT-1 (Dharmacon) as transfection reagent and siRNAs at a concentration of 25 nM. To reduce toxicity, the transfection medium containing siRNAs and reagent was removed after a 6-h incubation and replaced with fresh growth medium. Cells were incubated for additional 32 h prior to infection and further processing. In control transfections, siRNA-free siRNA buffer (Dharmacon) was added to the transfection medium instead of siRNAs.

### Fluorescence microscopy

To preserve cells for fluorescence microscopy, cells were fixed by a 10-min incubation in cold (−20°C) methanol or a 20-min incubation in 2-4% formaldehyde (with or without 0.025% glutaraldehyde) at room temperature. Formaldehyde-fixed cells (but not methanol-fixed cells) were then permeabilized for 15 min with 0.2% Triton X-100 in Dulbecco’s phosphate-buffered saline (DPBS; Gibco). Subsequently, the cells were incubated for 20 min in blocking solution (2-3% BSA in DPBS), prior to a 1-h incubation with primary antibodies diluted in blocking solution. Primary antibodies used included rabbit-anti-Slc1 (1:400; (Chen et al., 2014)), mouse-anti-FLAG (1:250-1:500; Sigma-Aldrich, F1804 and F3165), rat-anti-HA (1:500; Sigma-Aldrich, 11867423001), rabbit-anti-IPAM (1:100-1:200; gift from Daniel Rockey (Oregon State University)), rabbit-anti-CT222 (1:100; gift from Ted Hackstadt (NIH/NIAID, Rocky Mountain Laboratories), (Mital et al., 2010)), mouse-anti-phospho(Tyr416)-Src (1:100; Sigma-Aldrich, 05-677), and mouse-anti-α-tubulin (1:250-1:1000; Sigma-Aldrich, T5168). Following incubation with primary antibodies, cells were washed thrice with DPBS, incubated for 1 h with AlexaFluor (488, 555, or 647) -labeled secondary antibodies (Invitrogen) diluted in blocking solution (1:250-1:1000), and then washed again thrice with DPBS. When indicated, DNA was stained with 2-10 µg/ml Hoechst 33258 or 33342 (Invitrogen), during or after the incubation with secondary antibodies. In applications in which cells were grown and stained on glass coverslips, the coverslips were transferred to microscope slides and embedded in mowiol (18 ml 0.13 M Tris/HCl (pH 8.5), 2.4 g polyvinyl alcohol, 6 g glycerin, 0.01% p-phenylenediamine-dihydrochloride) or ProLong Glass Antifade Mountant (Invitrogen). Images were taken with various microscopy systems, including an epifluorescence microscope (Zeiss Axio Observer.Z1), inverted confocal microscopes (Zeiss LSM 780, Leica SP8), and high-content imaging platforms (ImageXpress Micro XL system (Molecular Devices), Cellomics ArrayScan VTI HCS (Thermo-Fisher-Scientific)).

### Co-immunoprecipitation

HeLa cells were infected with the indicated *Chlamydia* strains and when indicated transfected with EGFP-Rab expression plasmids at 1 hpi (see above). At 26 hpi, cells were lysed in Pierce IP lysis buffer supplemented with Pierce Protease and Phosphatase Inhibitor (Thermo-Fisher-Scientific). For improved extraction of CpoS variants, 0.2% sodium deoxycholate was added to the lysis buffer. Lysates were homogenized by passing them through a needle (0.33 x 12 mm) and were cleared by centrifugation (20000 x g, 4°C, 20 min). For precipitation of FLAG-tagged proteins, ANTI-FLAG M2 magnetic beads (Sigma-Aldrich) were incubated with the lysate (500-800 µl, 0.7-1.0 mg/ml protein) for 6 h at 4°C and were then washed four times with Pierce IP lysis buffer. Proteins were eluted with 0.1 M glycine/HCl (pH 3.0) and neutralized by addition of 1/5 volume of 0.5 M Tris/HCl, 1.5 M NaCl (pH 7.4). For the Co-IP of IPAM, proteins were instead eluted by boiling (10 min, 95°C) of the beads in NuPAGE LDS sample buffer (Thermo-Fisher-Scientific). For precipitation of EGFP-fusion proteins, GFP-Trap_MA magnetic beads (Chromotek) were incubated with lysate (1350 µl, 0.6 mg/ml protein; diluted in TBS/EDTA buffer (10 mM Tris/HCl, 150 mM NaCl, 0.5 mM EDTA pH 7.5)) for 5 h at 4°C and were then washed four times with TBS/EDTA buffer. Proteins were eluted from the beads by boiling (10 min, 95°C) in NuPAGE LDS sample buffer (Thermo-Fisher-Scientific). Aliquots from the initial lysate (lysate fraction), lysate after incubation with beads (unbound fraction), and the buffer used for the last wash step (wash fraction) were also collected. Samples were stored at −20°C until analysis by western blotting.

### Western blot analysis

For western blot analysis, protein extracts were generated by lysing cells in boiling 1% SDS buffer (50 mM Tris/HCl (pH 7.5), 1% SDS, 0.1 M NaCl), as previously described (Sixt et al., 2017), or were obtained from Co-IP experiments (see above). Protein samples were quantified using a BCA protein assay (Pierce) and adjusted to equal protein content by dilution with 1% SDS buffer (except for Co-IP samples). Samples were then mixed either with a commercial loading buffer (NuPAGE LDS sample buffer containing NuPAGE reducing agent (Thermo-Fisher-Scientific)) or a self-made loading buffer (final concentration: 50 mM Tris (pH 6.8), 10% glycerol, 2% SDS, 50 mM DTT, traces of bromophenol blue). Samples were denatured (10 min, 95-100°C), before separation by SDS PAGE using commercial gels (NuPAGE Novex 4-12% Bis-Tris gels (Thermo-Fisher-Scientific) or Mini-PROTEAN TGX 4-20% gels (Bio-Rad)) and transfer to 0.2 µm nitrocellulose membranes (Bio-Rad). Membranes were blocked for 1 h in blocking buffer (5% milk/TBST or 3% BSA/TBST) and incubated o/n at 4°C with primary antibody (diluted in blocking solution). Membranes were then washed three times for at least 15 min with TBST, followed by incubation with secondary antibodies (diluted in blocking solution) and three additional wash steps. Primary antibodies used included: rabbit-anti-CpoS (1:200; (Sixt et al., 2017)), mouse-anti-FLAG (1:1000; Sigma-Aldrich, F3165), rabbit anti-FLAG (1:1000; Sigma-Aldrich, F7425), rabbit-anti-Slc1 (1:1000; (Chen et al., 2014)), rabbit-anti-GFP (1:1000; Cell Signaling, 2956), rabbit-anti-MYC (1:1000; Cell Signaling, 2278), rabbit-anti-Rab35 (1:1000; Cell Signaling, 9690; or Proteintech, 11329-2-AP), mouse-anti-β-actin (1:2000; Cell Signaling, 3700) and HRP-conjugated mouse-anti-β-actin (1:50000; Abcam, ab49900). HRP-conjugated secondary antibodies (Southern Biotech or Thermo-Fisher-Scientific) were used at a dilution of 1:10000-1:50000. Membranes were incubated for 1 min with HRP substrate (ECL Prime Western Blotting Detection Reagent (GE Healthcare) or SuperSignal West Pico PLUS Chemiluminescent Substrate (Thermo-Fisher-Scientific)) and chemiluminescent signals were recorded with an Image Quant LAS4000 imaging system (GE Healthcare) or an Amersham Imager 680RGB (GE Healthcare). Membranes were stripped with Restore Plus western blot stripping buffer (Thermo-Fisher-Scientific) and blocked before detection of additional targets. For the quantitative analysis of CpoS and Rab35 expression, band intensities were quantified using software Image Quant TL (GE Healthcare). Expression levels of CpoS and Rab35 were normalized to the expression of the *Chlamydia* protein Slc1 and host β-actin, respectively.

### Quantification and statistical analysis

Where applicable, quantification and normalization of data was conducted as described above and/or in the respective figure legends. Pooled data are presented as mean±SD. Information regarding replication is indicated in the figure legends, where “n” denotes the number of independent biological experiments made. Statistical analysis was conducted using the software GraphPad Prism 8 using the statistical tests indicated in the figure legends (*, p<0.05; **, p<0.01; *** p<0.001; ns, not significant).

## DATA AVAILABILITY

This study includes no data deposited in external repositories.

## ACKNOWLEDGEMENTS

We acknowledge the Biochemical Imaging Center at Umeå University and the Swedish National Microscopy Infrastructure NMI (VR-RFI 2019-00217), the Chemical Biology Consortium Sweden, as well as the Duke University shared facilities, for providing assistance with microscopy and high-content imaging. We are grateful to Ted Hackstadt (RML/NIAID), Daniel Rockey (Oregon State University), Marci Scidmore (formerly Cornell University), Isabelle Dérre (University of Virginia), Derek J. Fisher (Southern Illinois University), and Kevin Hybiske (University of Washington) for sharing plasmids and reagents. BSS and work performed at Umeå University was supported by the European Union’s Seventh Framework Program (PIOF-GA-2013-626116) and the Swedish Research Council Vetenskapsrådet [2018-02286 to BSS and 2016-06598 to the Laboratory for Molecular Infection Medicine Sweden]. RHV and work performed at Duke University was supported by NIH awards AI100759 and AI134891. GK is supported by the Ligue contre le Cancer (équipe labellisée); Agence National de la Recherche (ANR) – Projets blancs; AMMICa US23/CNRS UMS3655; Association pour la recherche sur le cancer (ARC); Association “Ruban Rose”; Cancéropôle Ile-de-France; Fondation pour la Recherche Médicale (FRM); a donation by Elior; Equipex Onco-Pheno-Screen; European Joint Programme on Rare Diseases (EJPRD); Gustave Roussy Odyssea, the European Union Horizon 2020 Projects Oncobiome and Crimson; Fondation Carrefour; Institut National du Cancer (INCa); Inserm (HTE); Institut Universitaire de France; LabEx Immuno-Oncology (ANR-18-IDEX-0001); the Leducq Foundation; a Cancer Research ASPIRE Award from the Mark Foundation; the RHU Torino Lumière; Seerave Foundation; SIRIC Stratified Oncology Cell DNA Repair and Tumor Immune Elimination (SOCRATE); and SIRIC Cancer Research and Personalized Medicine (CARPEM). This study contributes to the IdEx Université de Paris ANR-18-IDEX-0001. OK receives funding from INCA and the DIM Elicit of the Ile de France.

## AUTHOR CONTRIBUTIONS

Conceptualization, KM, BSS; Investigation, KM, LHJ, LPJ, BSS; Formal analysis, KM, BSS; Visualization, KM, BSS; Writing – Original Draft, KM, BSS; Writing – Review & Editing, KM, BSS, OK, RHV, GK; Supervision, BSS, OK, RHV, GK; Funding acquisition, BSS, RHV, GK.

## CONFLICT OF INTEREST

The authors have no conflicts of interest to declare.

## SUPPORTING INFORMATION

**Figure S1.**
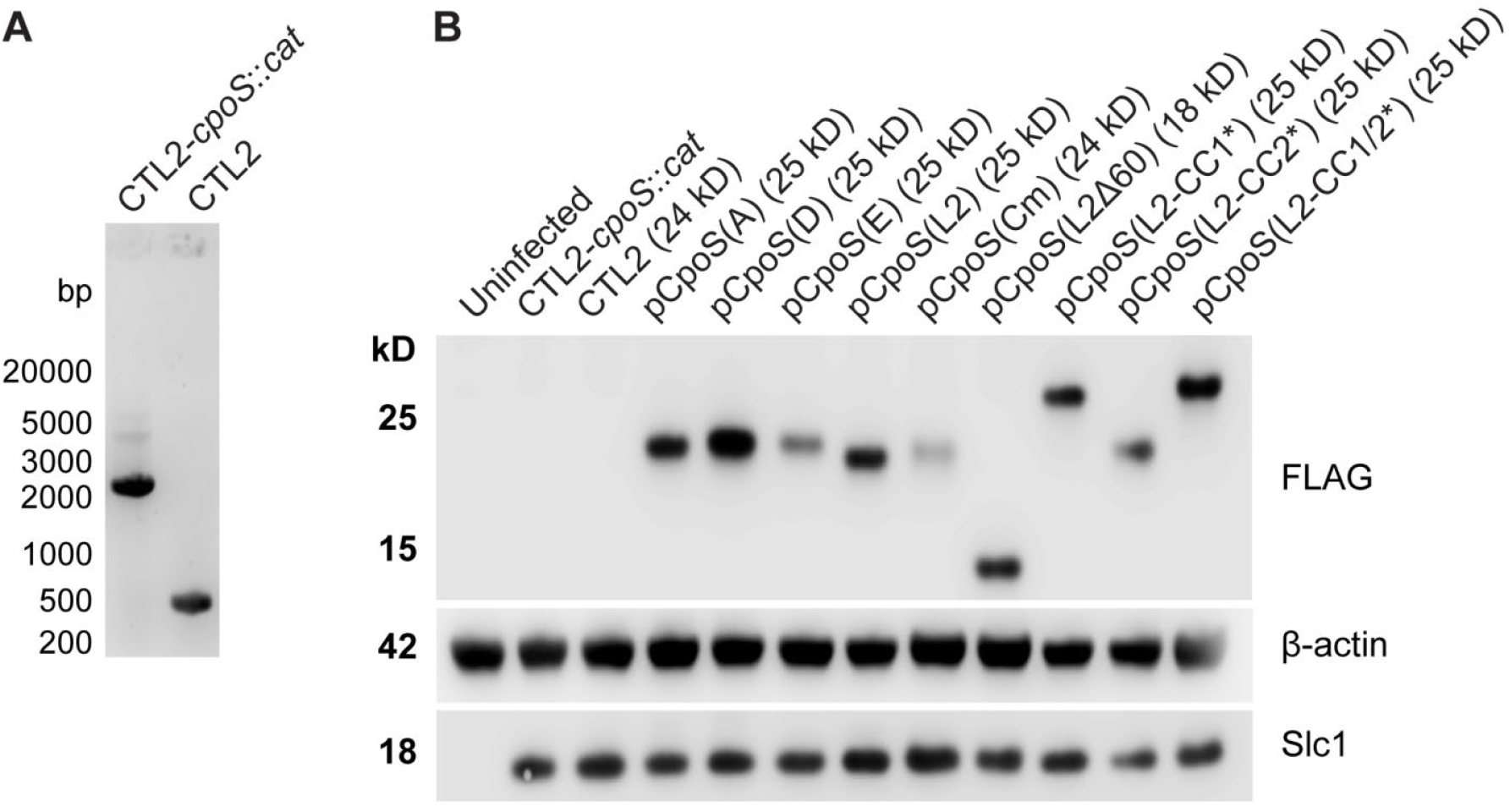
Insertional inactivation of *cpoS* with a *cat* resistance cassette and complementation with plasmid-expressed FLAG-tagged CpoS orthologs and variants. **(A)** PCR confirmation of insertional disruption of *cpoS* (CTL0481). Primers spanning the TargeTron insertion site amplify a band of 539 bp if the *cpoS* gene is intact and a band of 2428 bp if the intron insertion occurred in *cpoS* and disrupted the native locus. **(B)** Western blot analysis of CpoS-FLAG expression in HeLa cells infected for 32 h with the indicated strains (MOI20). The calculated molecular weight of each construct is indicated in parentheses. Deviations from the predicted size may reflect CpoS structure and structural changes introduced by the modifications.

**Figure S2.**
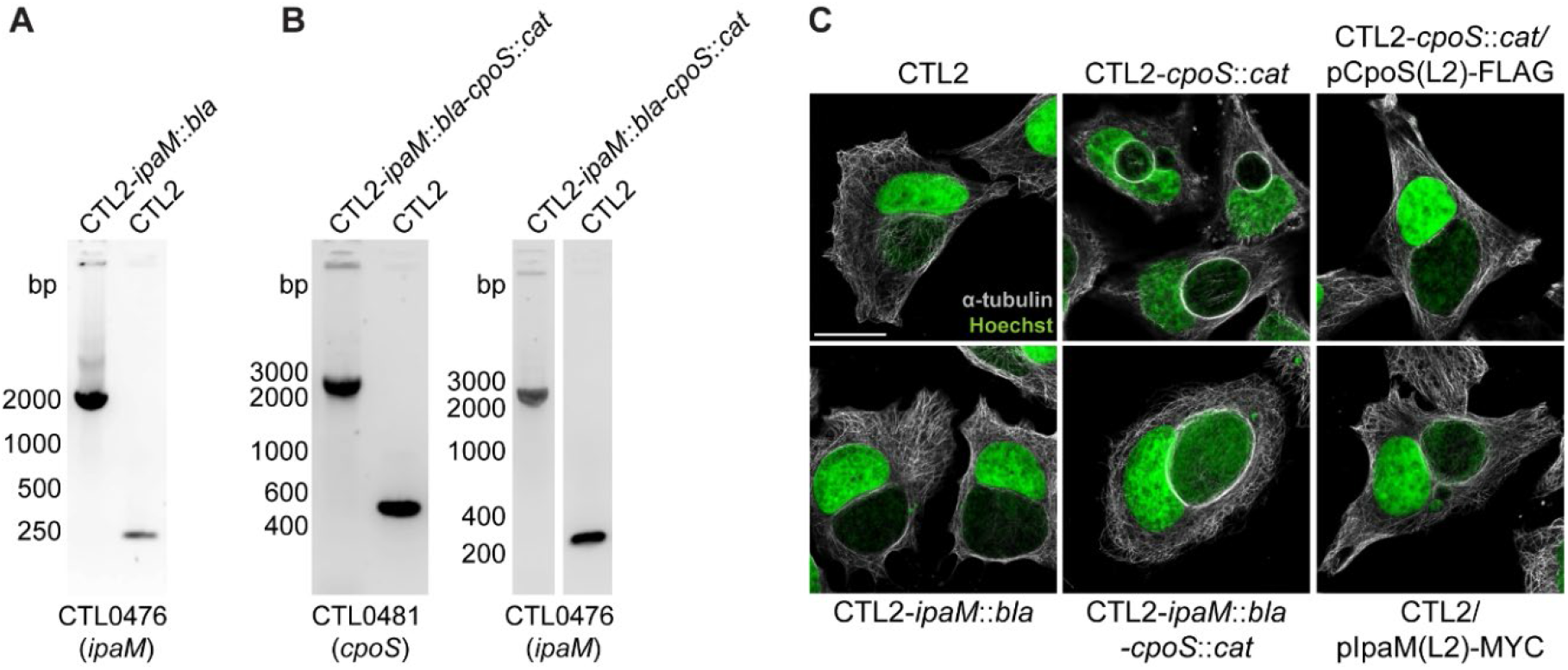
Insertional inactivation of *ipaM* and effects of IPAM-deficiency on microtubule architecture. **(A-B)** PCR confirmation of insertional disruption of *ipaM* (CTL0476) in the IPAM-deficient strain (CTL2-*ipaM*::*bla*) (A) or of *ipaM* (CTL0476) and *cpoS* (CTL0481) in the IPAM/CpoS double-deficient strain (CTL2-*ipaM*::*bla*-*cpoS*::*cat*) (B). Primers spanning the TargeTron insertion site amplify a band of 250 bp (CTL0476) or 539 bp (CTL0481) if the gene is intact and a band of 2331 bp (CTL0476) or 2428 bp (CTL0481) if the intron insertion occurred in the gene and disrupted the native locus. **(C)** Architecture of the microtubule cytoskeleton in infected HeLa cells (MOI10, 24 hpi, confocal, scale=20 µM).

**Figure S3.**
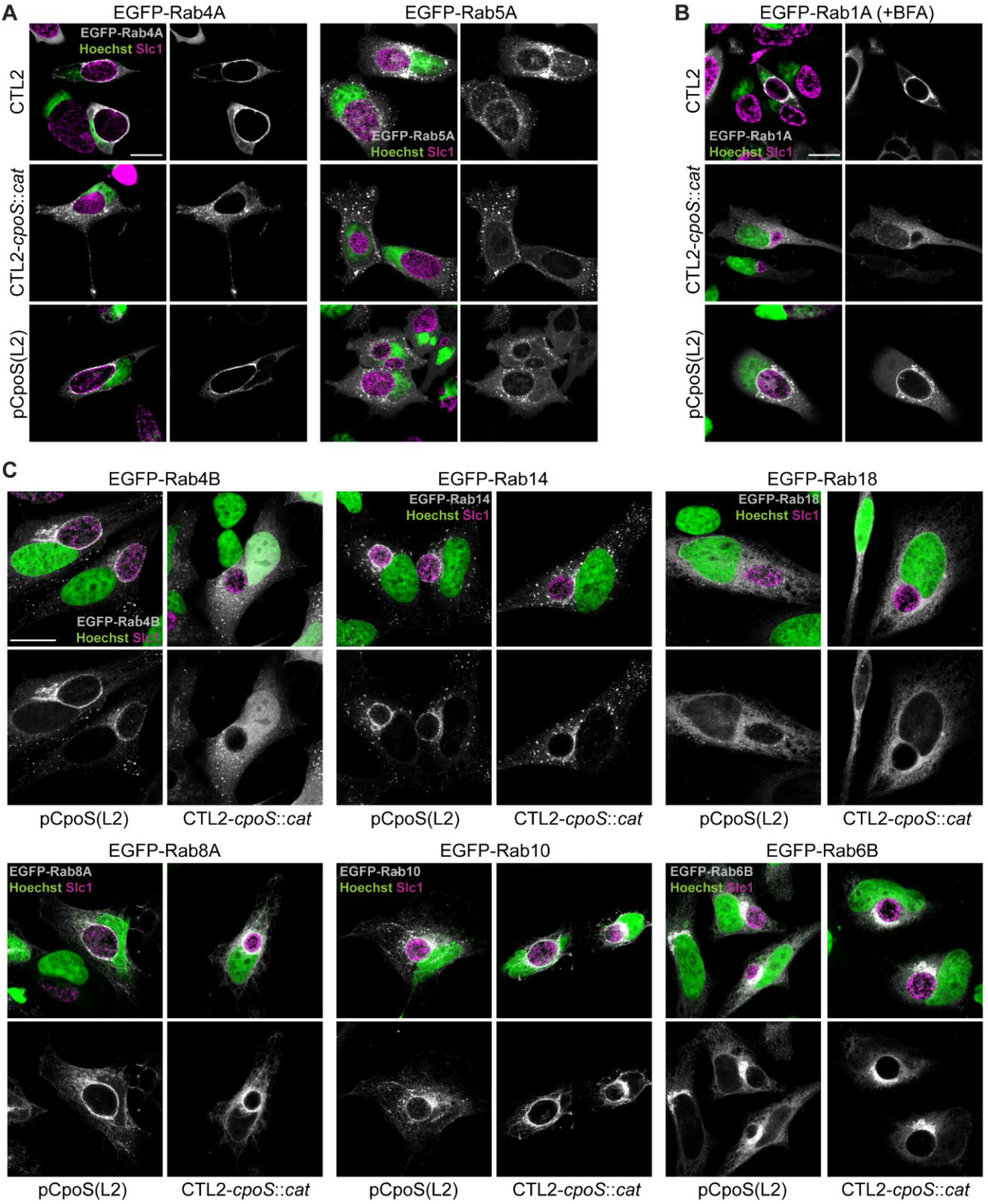
CpoS mediates recruitment of a diverse set of Rab proteins. **(A-C)** Localization of EGFP-Rab fusion proteins in infected HeLa cells (MOI5, 24 hpi, confocal, scale=20 µm). When indicated, BFA (3 µg/ml) was added at 12 hpi.

**Figure S4.**
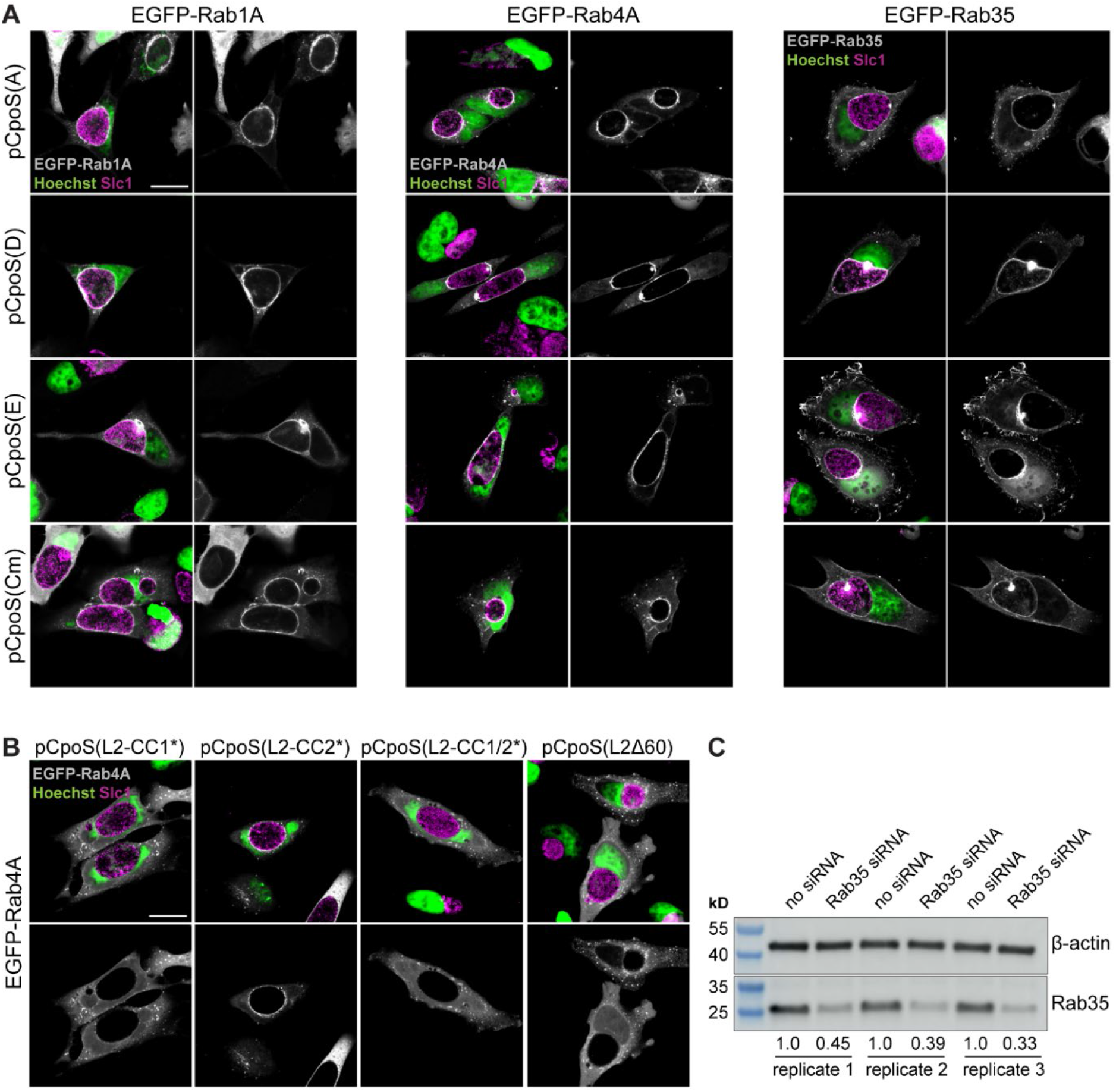
Differential abilities of CpoS orthologs and variants to mediate Rab recruitment, and efficiency of siRNA-mediated depletion of Rab35. **(A-B)** Localization of EGFP-Rab fusion proteins in infected HeLa (MOI5, 24 hpi, confocal, scale=20 µm). **(C)** Western blot analysis confirming depletion of Rab35 in A2EN cells treated with Rab35-specific siRNAs. Protein samples were generated at 46 h post transfection in three independent replicates. Relative Rab35 expression levels are indicated below the bands.

**Table S1.** Chlamydia strains used in this study.

| Strain name | Short name | Description | CpoS variant expressed |
| --- | --- | --- | --- |
| CTL2 | CTL2 | *C. trachomatis* strain L2/434/Bu | Endogenous CpoS(L2) |
| CTL2-*cpoS*::*cat* | CTL2-*cpoS*::*cat* | Derived from CTL2, *cat* insertion in *cpoS* | CpoS(L2) truncated after aa 92 |
| CTL2-*ipaM*::*bla* | CTL2-*ipaM*::*bla* | Derived from CTL2, *bla* insertion in *ipaM* | Endogenous CpoS(L2) |
| CTL2-*ipaM*::*bla-cpoS*::*cat* | CTL2-*ipaM*::*bla-cpoS*::*cat* | Derived from CTL2, *bla* insertion in *ipaM*, *cat* insertion in *cpoS* | CpoS(L2) truncated after aa 92 |
| CTL2-*cpoS*::*cat* / pCpoS(L2)-FLAG | pCpoS(L2) | Derived from CTL2-*cpoS*::*cat*, transformed with plasmid enabling expression of FLAG-tagged CpoS from *C. trachomatis* strain L2/434/Bu (CTL0481) | Plasmid-expressed CpoS(L2)-FLAG |
| CTL2-*cpoS*::*cat* / pCpoS(A)-FLAG | pCpoS(A) | Derived from CTL2-*cpoS*::*cat*, transformed with plasmid enabling expression of FLAG-tagged CpoS from *C. trachomatis* strain A/HAR-13 (CTA0251) | Plasmid-expressed CpoS(A)-FLAG |
| CTL2-*cpoS*::*cat* / pCpoS(D)-FLAG | pCpoS(D) | Derived from CTL2-*cpoS*::*cat*, transformed with plasmid enabling expression of FLAG-tagged CpoS from *C. trachomatis* strain D/UW-3/CX (CT229) | Plasmid-expressed CpoS(D)-FLAG |
| CTL2-*cpoS*::*cat* / pCpoS(E)-FLAG | pCpoS(E) | Derived from CTL2-*cpoS*::*cat*, transformed with plasmid enabling expression of FLAG-tagged CpoS from *C. trachomatis* strain E/Bour (BOUR00241) | Plasmid-expressed CpoS(E)-FLAG |
| CTL2-*cpoS*::*cat* / pCpoS(Cm)-FLAG | pCpoS(Cm) | Derived from CTL2-*cpoS*::*cat*, transformed with plasmid enabling expression of FLAG-tagged CpoS from *C. muridarum* strain Nigg (TC0500) | Plasmid-expressed CpoS(Cm)-FLAG |
| CTL2-*cpoS*::*cat* / pCpoS(L2Δ60)-FLAG | pCpoS(L2Δ60) | Derived from CTL2-*cpoS*::*cat*, transformed with plasmid enabling expression of FLAG-tagged CpoS from *C. trachomatis* strain L2/434/Bu (CTL0481) with a 60 aa C-terminal deletion | Plasmid-expressed CpoS(L2Δ60)-FLAG |
| CTL2-*cpoS*::*cat* / pCpoS(L2-CC1*)-FLAG | pCpoS(L2-CC1*) | Derived from CTL2-*cpoS*::*cat*, transformed with plasmid enabling expression of FLAG-tagged CpoS from *C. trachomatis* strain L2/434/Bu (CTL0481) with two point mutations in CC1 (I99P + L113P) | Plasmid-expressed CpoS(L2-CC1*)-FLAG (=CpoS(L2-I99P-L113P)-FLAG) |
| CTL2-*cpoS*::*cat* / pCpoS(L2-CC2*)-FLAG | pCpoS(L2-CC2*) | Derived from CTL2-*cpoS*::*cat*, transformed with plasmid enabling expression of FLAG-tagged CpoS from *C. trachomatis* strain L2/434/Bu (CTL0481) with two point mutations in CC2 (N181P + L188P) | Plasmid-expressed CpoS(L2-CC2*)-FLAG (=CpoS(L2-N181P-L188P)-FLAG) |
| CTL2-*cpoS*::*cat* / pCpoS(L2-CC1/2*)-FLAG | pCpoS(L2-CC1/2*) | Derived from CTL2-*cpoS*::*cat*, transformed with plasmid enabling expression of FLAG-tagged CpoS from *C. trachomatis* strain L2/434/Bu (CTL0481) with two point mutations in CC1 (I99P + L113P) and two point mutations in CC2 (N181P + L188P) | Plasmid-expressed CpoS(L2-CC1/2*)-FLAG (=CpoS(L2-I99P-L113P-N181P-L188P)-FLAG) |
| CTL2 / pIPAM(L2)-MYC | pIPAM(L2)-MYC | Derived from CTL2, transformed with plasmid enabling expression of MYC-tagged IPAM from *C. trachomatis* strain L2/434/Bu (CTL0476) | Endogenous CpoS(L2) |
| CTL2 / p2TK2-SW2-mCherry | n/a | Derived from CTL2, transformed with plasmid enabling expression of mCherry | Endogenous CpoS(L2) |
| CTL2-*cpoS*::*cat* / p2TK2-SW2-mCherry | n/a | Derived from CTL2-*cpoS*::*cat*, transformed with plasmid enabling expression of mCherry | CpoS(L2) truncated after aa 92 |
| CTL2 / pTL2-tetO-IncB-GFP11x7-FLAG | CTL2/plncB-GFP11 | Derived from CTL2, transformed with plasmid enabling expression of IncB tagged with seven repeats of GFP11 and a FLAG tag | Endogenous CpoS(L2) |
| CTL2-*cpoS*::*cat* / pTL2-tetO-IncB-GFP11x7-FLAG | CTL2-*cpoS*::*cat*/plncB-GFP11 | Derived from CTL2-*cpoS*::*cat*, transformed with plasmid enabling expression of IncB tagged with seven repeats of GFP11 and a FLAG tag | CpoS(L2) truncated after aa 92 |

**Table S2.**
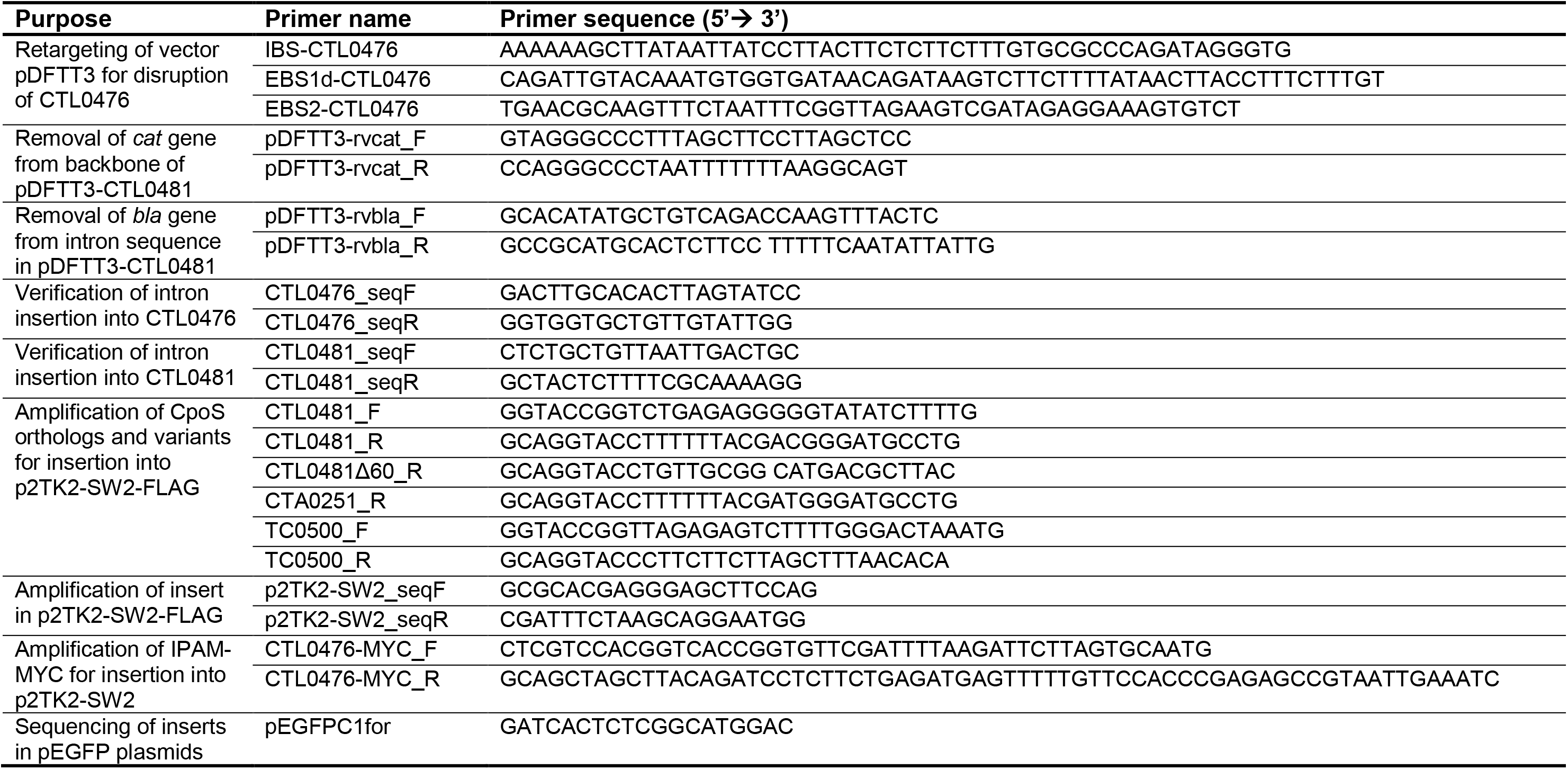
Primers used in this study.

**Table S3.** siRNAs used in this study.

| **Reagent** | **Vendor** | **Catalog #** |
| --- | --- | --- |
| siGENOME siRNA SMARTpool targeting human STING | Dharmacon | M-024333-00-0005 |
| siGENOME siRNA SMARTpool targeting human Rab1A | Dharmacon | M-008283-01-0005 |
| siGENOME siRNA SMARTpool targeting human Rab1B | Dharmacon | M-008958-01-0005 |
| siGENOME siRNA SMARTpool targeting human Rab2A | Dharmacon | M-010533-01-0005 |
| siGENOME siRNA SMARTpool targeting human Rab2B | Dharmacon | M-010370-00-0005 |
| siGENOME siRNA SMARTpool targeting human Rab4A | Dharmacon | M-008539-01-0005 |
| siGENOME siRNA SMARTpool targeting human Rab5A | Dharmacon | M-004009-00-0005 |
| siGENOME siRNA SMARTpool targeting human Rab8A | Dharmacon | M-003905-00-0005 |
| siGENOME siRNA SMARTpool targeting human Rab8B | Dharmacon | M-008744-01-0005 |
| siGENOME siRNA SMARTpool targeting human Rab35 | Dharmacon | M-009781-00-0005 |

## Notes

### Competing Interest Statement

The authors have declared no competing interest.

